# Identification of candidate biomarkers and pathways associated with type 1 diabetes mellitus using bioinformatics analysis

**DOI:** 10.1101/2021.06.08.447531

**Authors:** Basavaraj Vastrad, Chanabasayya Vastrad

## Abstract

Type 1 diabetes mellitus (T1DM) is a metabolic disorder for which the underlying molecular mechanisms remain largely unclear. This investigation aimed to elucidate essential candidate genes and pathways in T1DM by integrated bioinformatics analysis. In this study, differentially expressed genes (DEGs) were analyzed using DESeq2 of R package from GSE162689 of the Gene Expression Omnibus (GEO). Gene ontology (GO) enrichment analysis, REACTOME pathway enrichment analysis, and construction and analysis of protein-protein interaction (PPI) network, modules, miRNA-hub gene regulatory network and TF-hub gene regulatory network, and validation of hub genes were then performed. A total of 952 DEGs (477 up regulated and 475 down regulated genes) were identified in T1DM. GO and REACTOME enrichment result results showed that DEGs mainly enriched in multicellular organism development, detection of stimulus, diseases of signal transduction by growth factor receptors and second messengers, and olfactory signaling pathway. The top hub genes such as MYC, EGFR, LNX1, YBX1, HSP90AA1, ESR1, FN1, TK1, ANLN and SMAD9 were screened out as the critical genes among the DEGs from the PPI network, modules, miRNA-hub gene regulatory network and TF-hub gene regulatory network. Receiver operating characteristic curve (ROC) analysis and RT-PCR confirmed that these genes were significantly associated with T1DM. In conclusion, the identified DEGs, particularly the hub genes, strengthen the understanding of the advancement and progression of T1DM, and certain genes might be used as candidate target molecules to diagnose, monitor and treat T1DM.

## Introduction

Type 1 diabetes mellitus (T1DM) is chronic autoimmune diabetes characterized by autoimmune mediated destruction of pancreatic beta cells [1]. T1DM is most generally identified in children and adolescents [2]. Epidemiological studies have shown that the incidence of T1DM has been increasing by 2% to 5% globally [3]. T1DM is a complex disease affected by numerous environmental factors, genetic factors and their interactions [4–5]. Several complication of T1DM such as cardiovascular disease [6], hypertension [7], diabetic retinopathy [8], diabetic nephropathy [9], diabetic neuropathy [10], obesity [11] and cognitive impairment [12]. Therefore, exploring the molecular mechanism of T1DM development and its associated biomarkers to improve the early diagnosis and treatment of T1DM.

Although the remarkable improvement is achieved in the treatment of T1DM is insulin therapy [13], the long-term survival rates of T1DM still remain low worldwide. One of the major reasons is that most patients with T1DM were diagnosed at advanced stages. It is crucial to find out new therapeutic targets and novel diagnostic biomarkers for the early diagnosis and timely treatment of T1DM. Therefore, it is still urgent to further explore the exact molecular mechanisms of the development of T1DM. At present, several genes and signaling pathway are identified; for example vitamin D receptor (VDR) [14], HLA-B and HLA-A [15], HLA-DQ [16], HLA DQB1, HLA DQA1 and HLA DRB1 [17], IDDM2 [18], CaMKII/NF-B/TG L 1 and PP L signaling L ay [19], Keap1/Nrf2 κ F-β AR-γ pathw signaling pathway [20], HIF-1/VEGF pathway [21], NLRP3 and NLRP1 inflammasomes signaling pathway [22] and NO/cGMP signaling pathway [23]. Therefore, it is of great practical significance to explore the genes and signaling pathways of T1DM on β islet cells.

High-throughput RNA sequencing platform for gene expression analysis have been increasingly recognized as approaches with significant clinical value in areas such as molecular diagnosis, prognostic prediction and identification of novel therapeutic targets [24]. RNA sequencing analysis has been widely used in various gene expression profiling studies examining disease pathogenesis in the last decade, and has revealed many differentially expressed genes (DEGs) associated in various pathways, biological processes, and molecular functions. We therefore used an expression profiling by high throughput sequencing dataset to investigate the pathogenesis of T1DM.

We downloaded expression profiling by high throughput sequencing dataset GSE162689 [25], from Gene Expression Omnibus database (GEO) (http://www.ncbi.nlm.nih.gov/geo/) [26], which contain gene expression data from T1DM samples and normal control samples. We then performed deep bioinformatics analysis, including identifying common differential expressed genes (DEGs), gene ontology (GO) enrichment analysis, REACTOME pathway enrichment analysis, and construction and analysis of protein-protein interaction (PPI) network, modules, miRNA-hub gene regulatory network and TF-hub gene regulatory network. The findings were further validated by receiver operating characteristic curve (ROC) analysis and RT-PCR. The aim of this study was to identify DEGs and important pathways, and to explore potential candidate biomarkers for the diagnosis, prognosis and therapeutic targets in T1DM.

## Materials and methods

### Data resources

Expression profiling by high throughput sequencing dataset GSE162689 [25] was obtained from GEO database. The dataset comprised total 59 samples, of which 27 were from T1DM samples and 32 were from normal control samples and was based on the GPL24014 Ion Torrent S5 XL (Homo sapiens).

### Identification of DEGs

Differentially expressed genes (DEGs) between T1DM and normal control samples were identified by using the DESeq2 package on R language software [27]. DEGs were considered when an adjusted P < 0.05, and a |log2 fold change| > 0.63 for up regulated genes and |log2 fold change| < -1.3 for down regulated genes. The adjusted P values, by employing Benjamini and Hochberg false discovery rate [28], were aimed to correct the occurrence of false positive results. The DEGs were presented in volcano plot and heat map drawn using a plotting tool ggplot2 and gplots based on the R language.

### GO and REACTOME pathway enrichment analysis of DEGs

One online tool, g:Profiler (http://biit.cs.ut.ee/gprofiler/) [29], was applied to carried out the functional annotation for DEGs. Gene Ontology (GO) (http://geneontology.org/) [30] generally perform enrichment analysis of genomes. And there are mainly biological processes (BP), cellular components (CC) and molecular functions (MF) in the GO enrichment analysis. REACTOME (https://reactome.org/) [31] is a comprehensive database of genomic, chemical, and systemic functional information. Therefore, g:Profiler was used to make enrichment analysis of GO and REACTOME. P < 0.05 was set as the cutoff criterion.

### Construction of the PPI network and module analysis

PPI network was established using the IntAct Molecular Interaction Database (https://www.ebi.ac.uk/intact/) [32]. To assess possible PPI correlations, previously identified DEGs were mapped to the IntAct database, followed by extraction of PPI pairs with a combined score >0.4. Cytoscape 3.8.2 software (www.cytoscape.org/) [33] was then employed to visualize the PPI network, and the Cytoscape plugin Network Analyzer was used to calculate the node degree [34], betweenness centrality [35], stress centrality [36] and closeness centrality [37] of each protein node. Specifically, nodes with a higher node degree, betweenness centrality, stress centrality and closeness centrality were likely to play a more vital role in maintaining the stability of the entire network. The PEWCC1 (http://apps.cytoscape.org/apps/PEWCC1) [38] plug-in was applied to analyze the modules in the PPI networks, with the default parameters (node score = 0.2, K-core≧2, and max depth = 100).

### MiRNA-hub gene regulatory network construction

The miRNAs targeting the T1DM related were predicted using the miRNet database (https://www.mirnet.ca/) [39], and those predicted by at least 14 databases (TarBase, miRTarBase, miRecords, miRanda, miR2Disease, HMDD, PhenomiR, SM2miR, PharmacomiR, EpimiR, starBase, TransmiR, ADmiRE, and TAM 2.0) were selected for constructing the miRNA-hub gene regulatory network by Cytoscape 3.8.2 software.

### TF-hub gene regulatory network construction

The TFs targeting the T1DM related were predicted using the NetworkAnalyst database (https://www.networkanalyst.ca/) [40], and those predicted by RegNetwork database was selected for constructing the TF-hub gene regulatory network by Cytoscape 3.8.2 software.

### Validation of hub genes by receiver operating characteristic curve (ROC) analysis

A ROC curve analysis is an approach for visualizing, organizing and selecting classifiers based on their achievement of hub genes. A diagnostic test was firstly performed in order to estimate the diagnostic value of hub genes in T1DM. ROC curves were obtained by plotting the sensitivity, against the specificity using the R package “pROC” [41]. Area under the curve (AUC) was used to measure the that the model had a favorable fitting effect.

### Detection of the mRNA expression of the hub genes by RT-PCR

Pancreatic beta MIN6 cells were maintained in Dulbecco’s minimal essential medium (DMEM) supplemented with 15 % fetal calf serum, 50 mg/l streptomycin and 75 mg/l penicillin sulphate. MIN6 cells treated with streptozotocin for T2DM and MIN6 cells untreated for normal control. All treated MIN6 cells and untreated MIN6 cells were incubated at 37 ◦ 5% CO_2_ in humidified incubator. Total RNA was extracted from the treated and untreated cells using TRI reagent (Sigma, USA) according to the manufacturer’s protocol. Reverse transcription kit (Thermo Fisher Scientific, Waltham, MA, USA) was used for Converting mRNA to cDNA. Expression levels of mRNAs were determined by RT-PCR using QuantStudio 7 Flex real-time PCR system (Thermo Fisher Scientific, Waltham, MA, USA). The protocol was set as follows: 50 °C for 2 min, 95 °C for 10L n, 40L ycles of 95 °C for 10 s, 60L for 30L All the samples were normalized to the -actin . The test was performed in corresponding expression of internal control β triplicate and the relative expression levels were calculated with the 2^-ΔΔCt^ method [42]. Table 1 is given for primer sequences used in the RT-PCR.

**Table 1.**
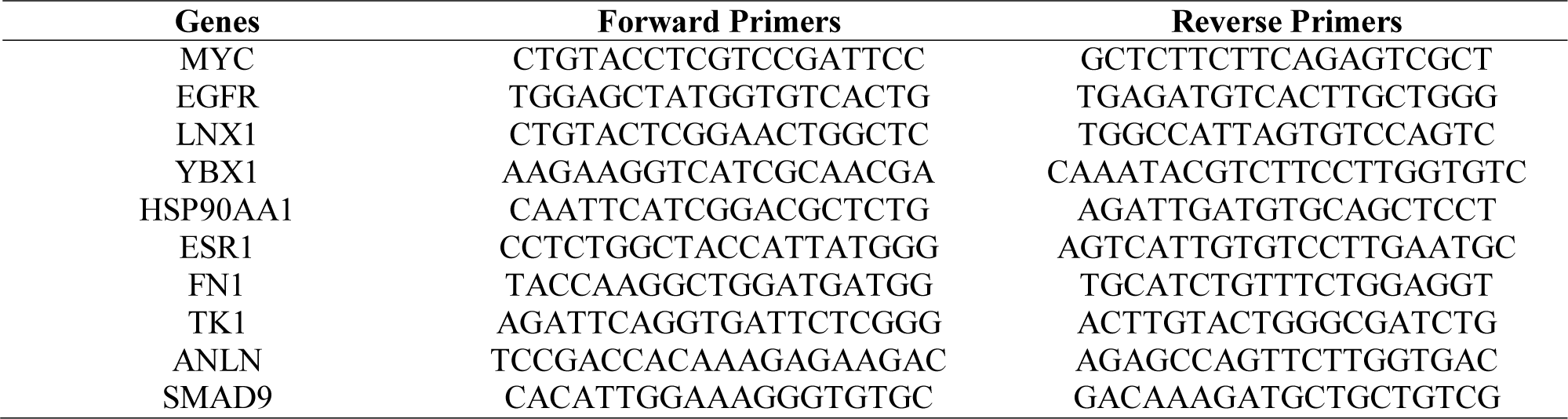
The sequences of primers for quantitative RT-PCR

## Results

### Identification of DEGs

On the basis of the cut off criteria, DEGs in GEO dataset was identified between T1DN and normal co L l samples (Table 2). There were 952 DEGs, including 477 up regulated and 475 down regulated genes in GSE162689 with the threshold of adjusted P < 0.05, and a |log2 fold change| > 0.63 for up regulated genes and |log2 fold change| < -1.3 for down regulated genes. The volcano plot of the distribution of DEGs in is shown in Fig. 1. The expression heat map of the DEGs is shown in Fig. 2.

**Fig. 1.**
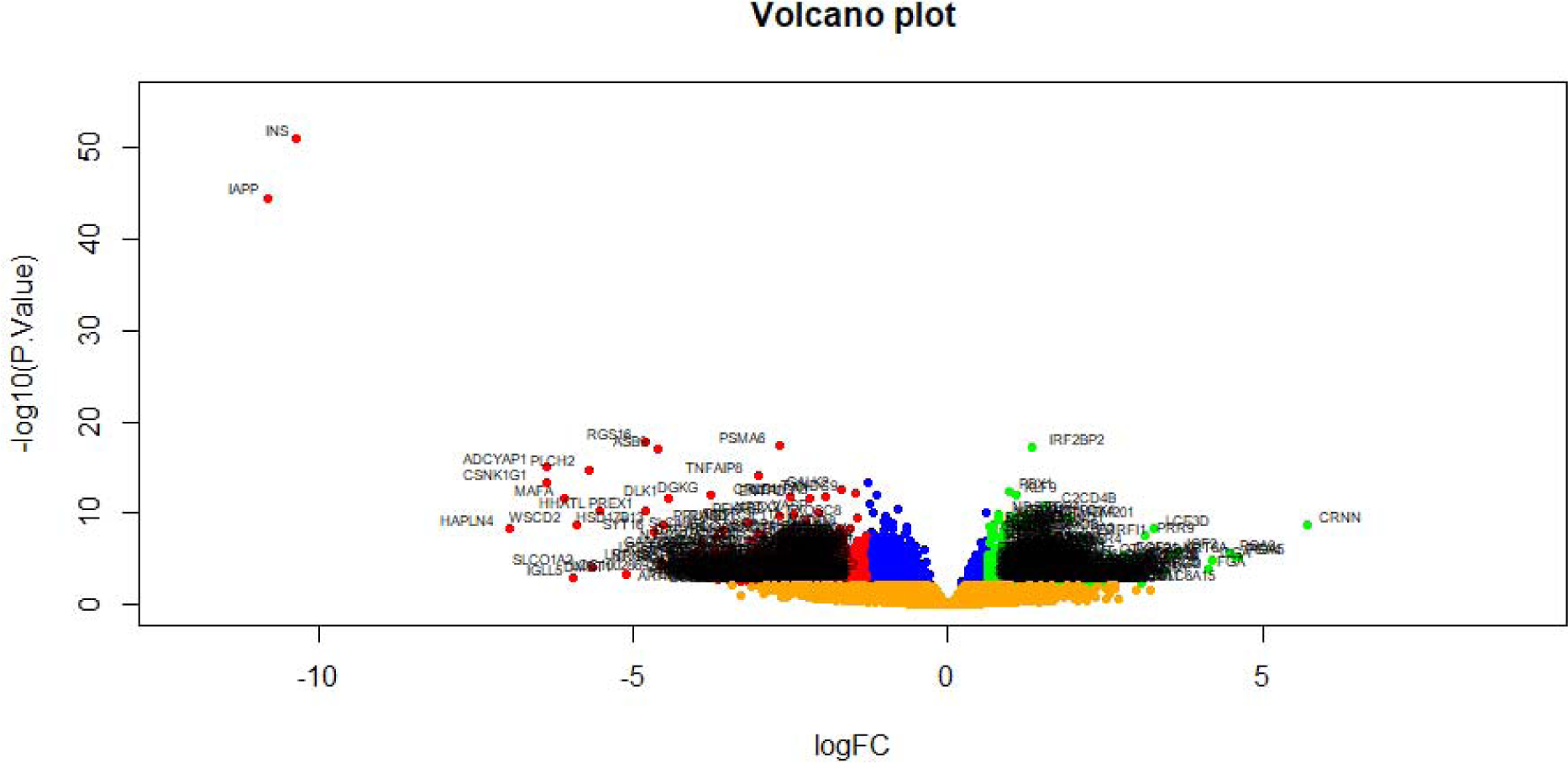
Volcano plot of differentially expressed genes. Genes with a significant change of more than two-fold were selected. Green dot represented up regulated significant genes and red dot represented down regulated significant genes.

**Fig. 2.**
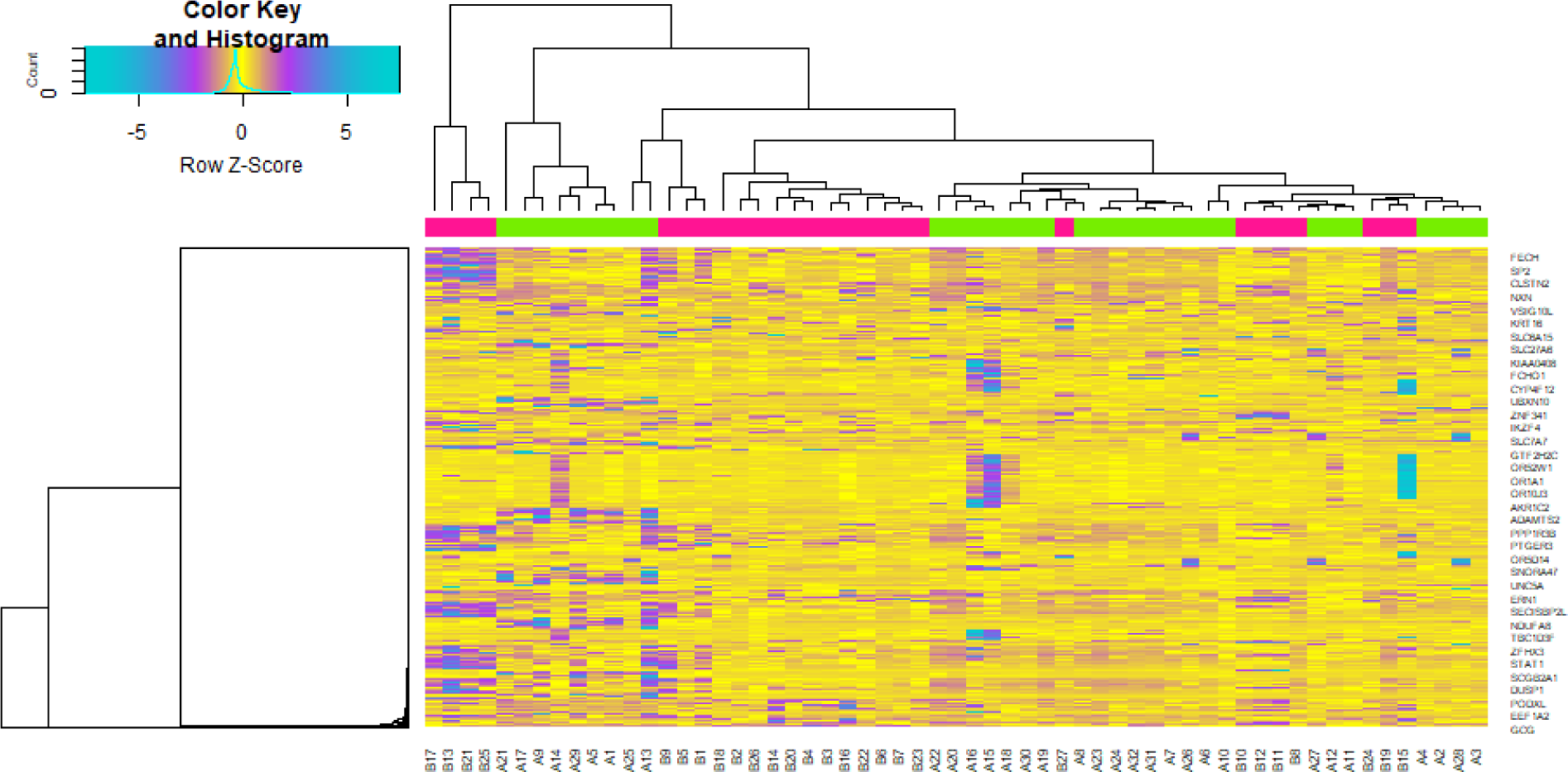
Heat map of differentially expressed genes. Legend on the top left indicate log fold change of genes. (A1 – A32= normal control samples; B1 – B27 = T1DM samples)

**Table 2.**
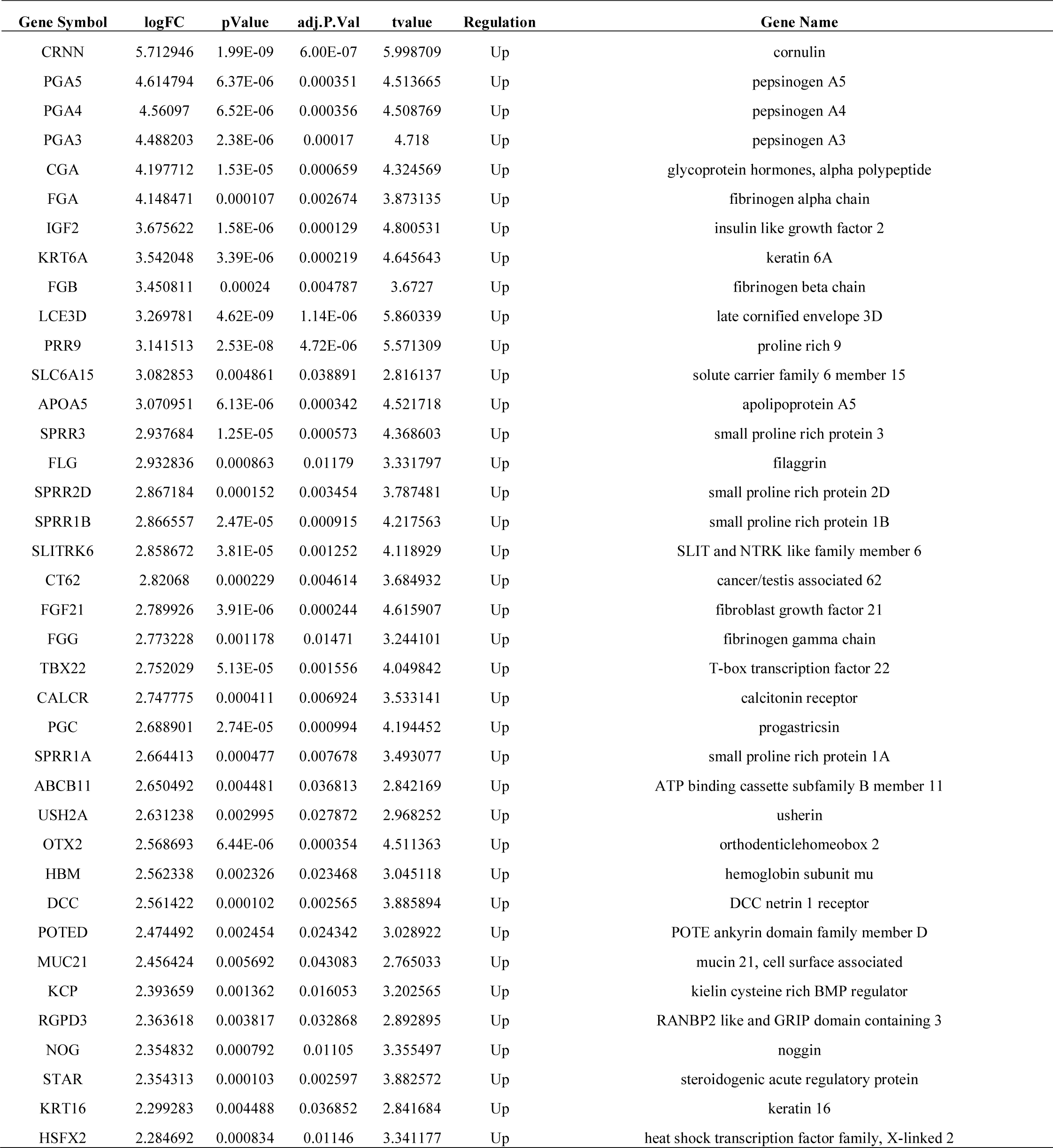
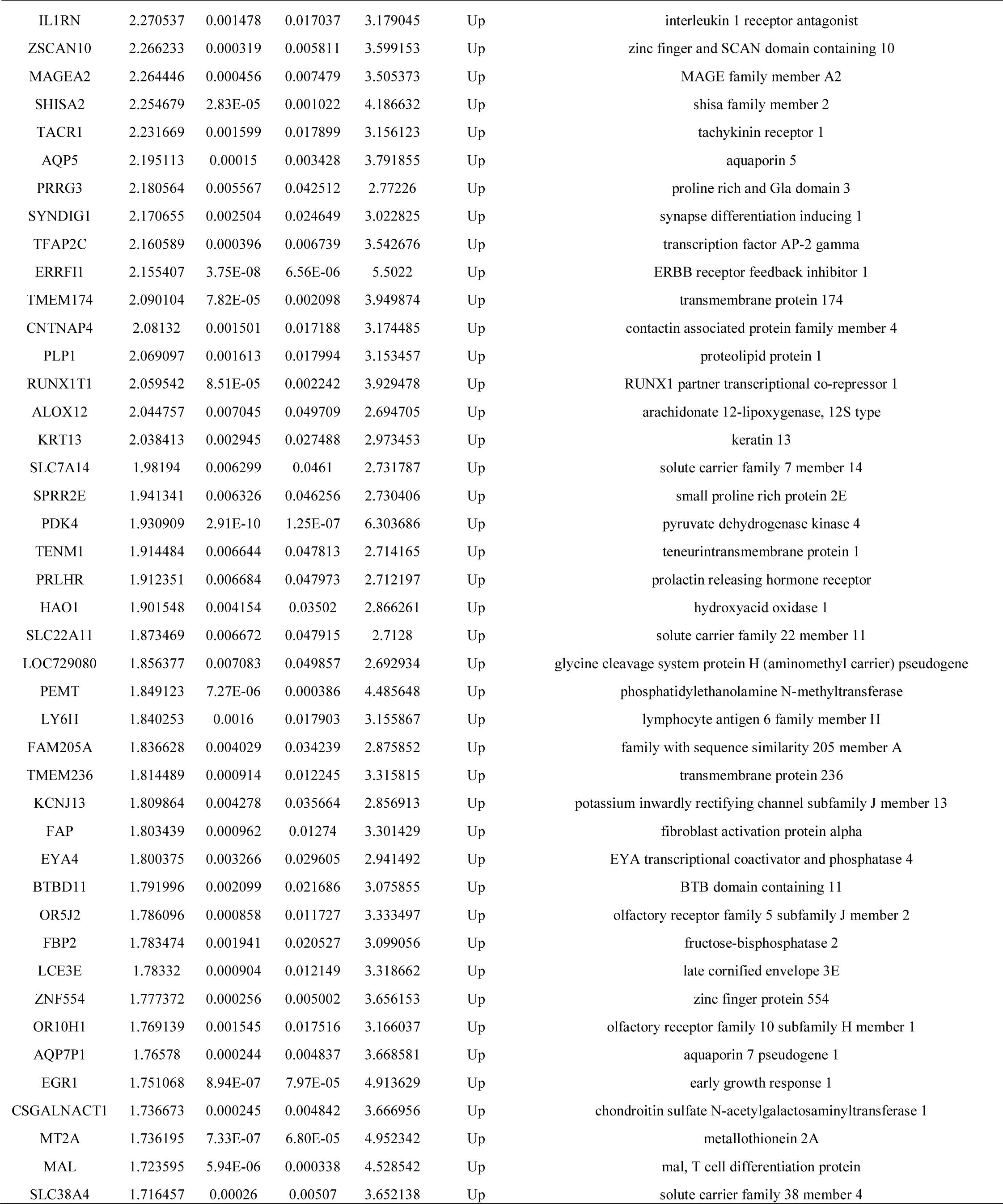
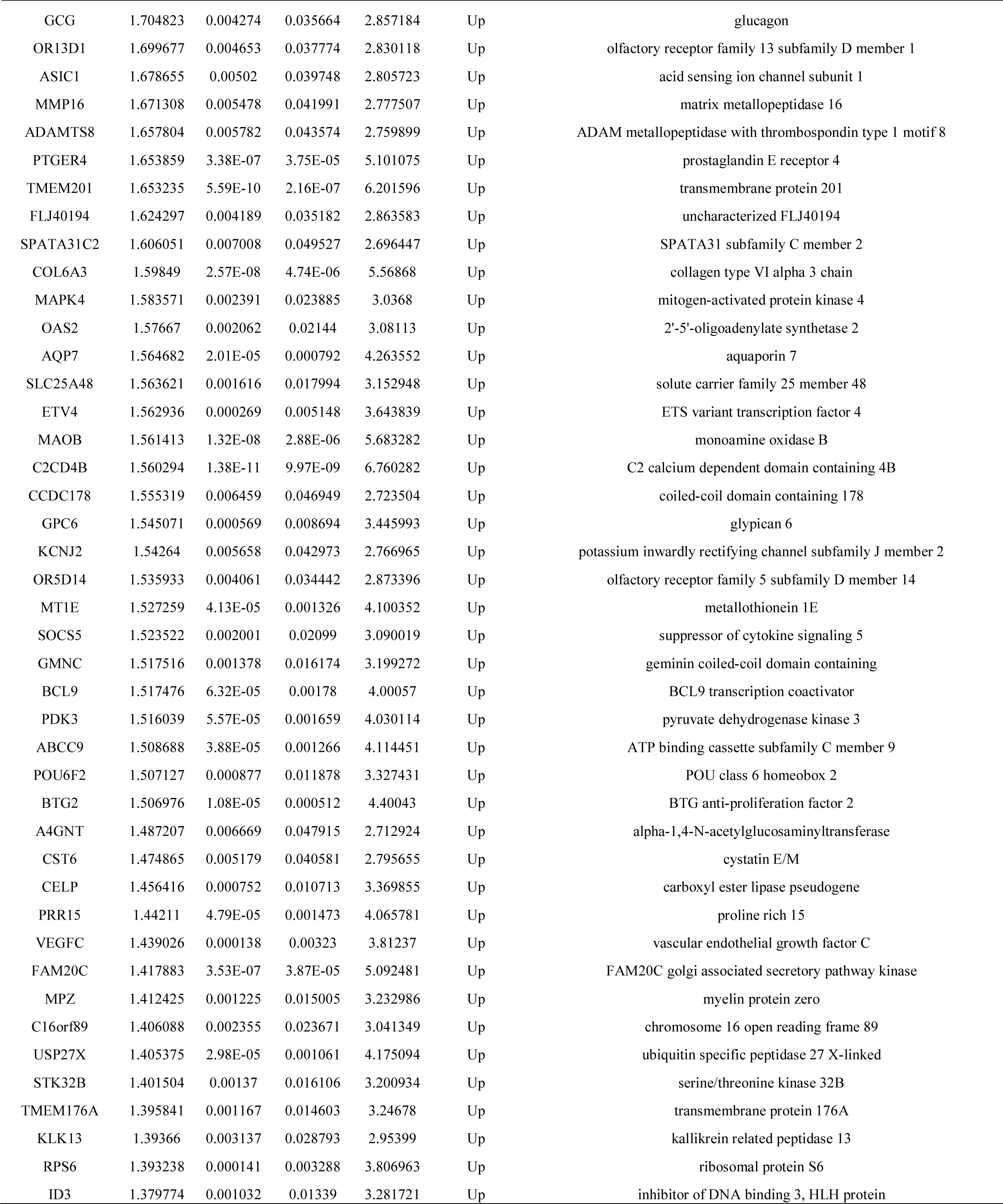
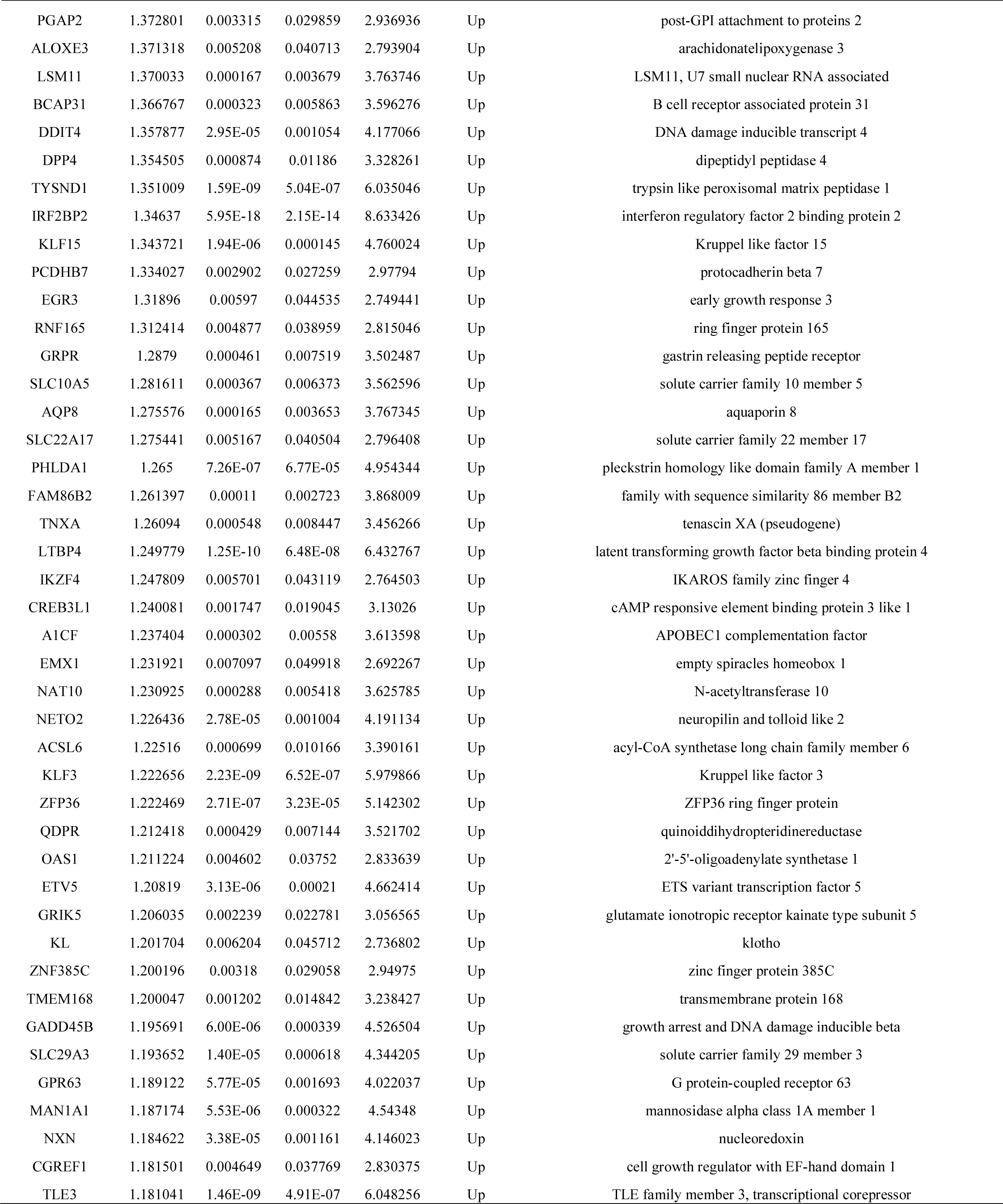
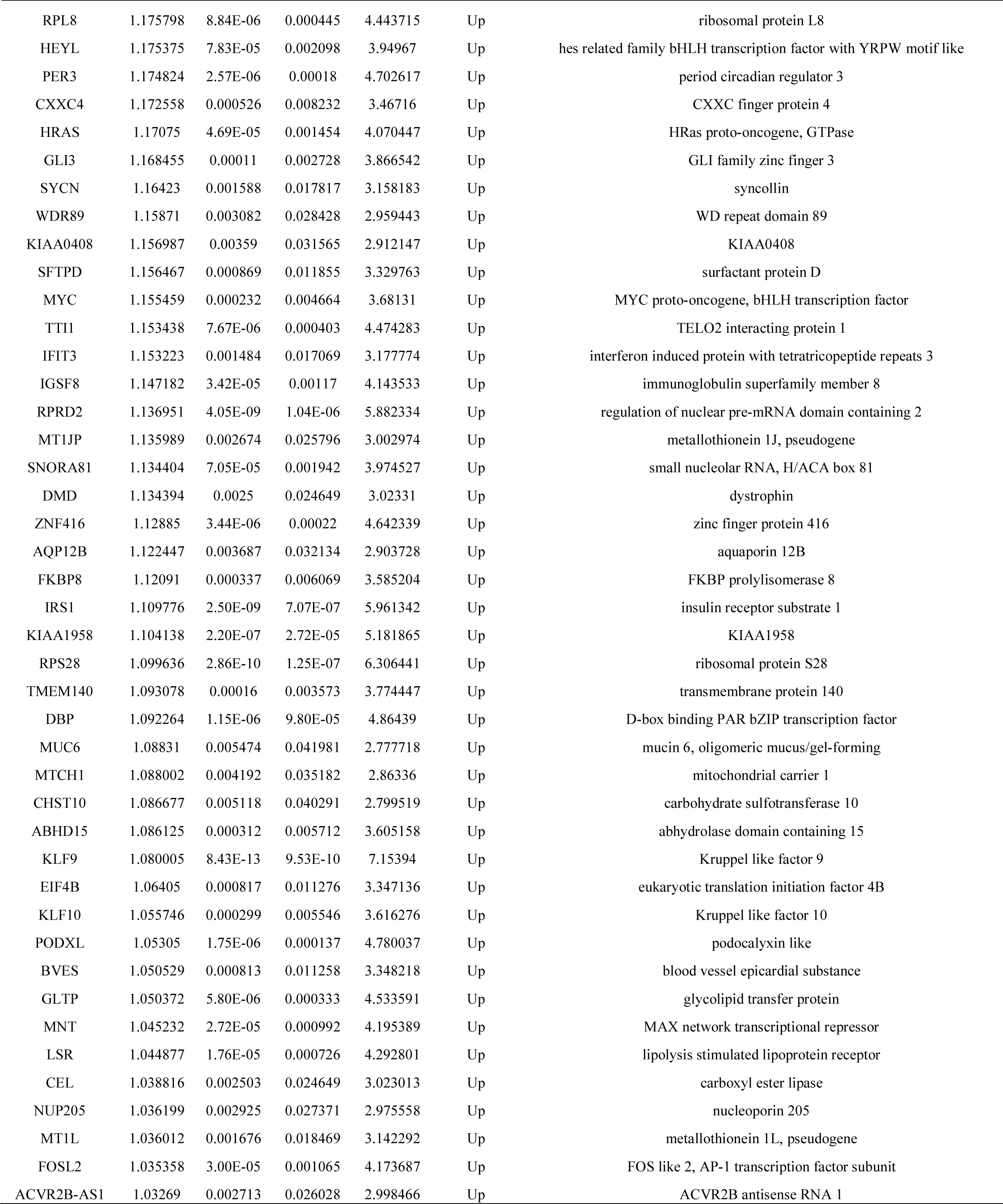
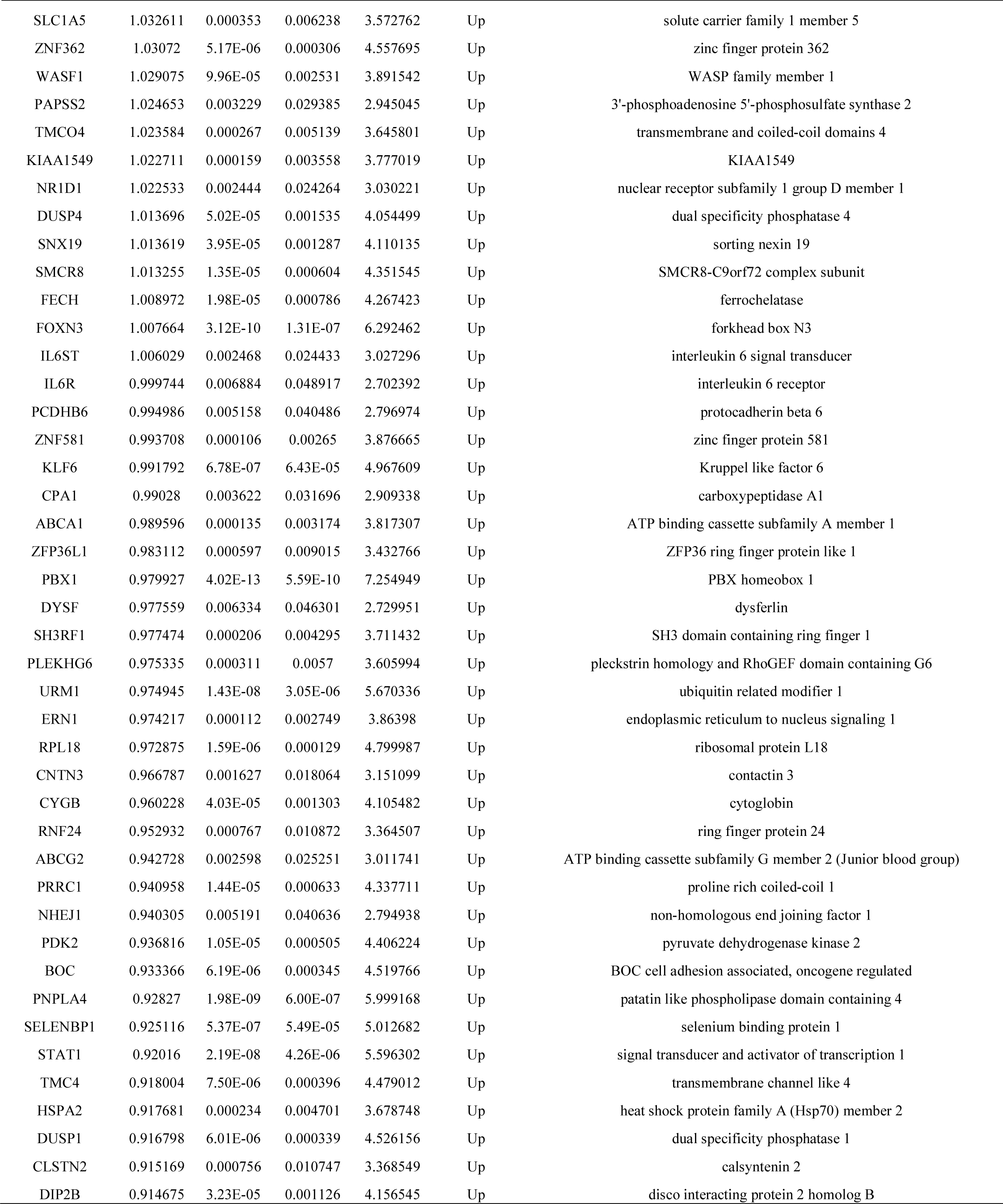
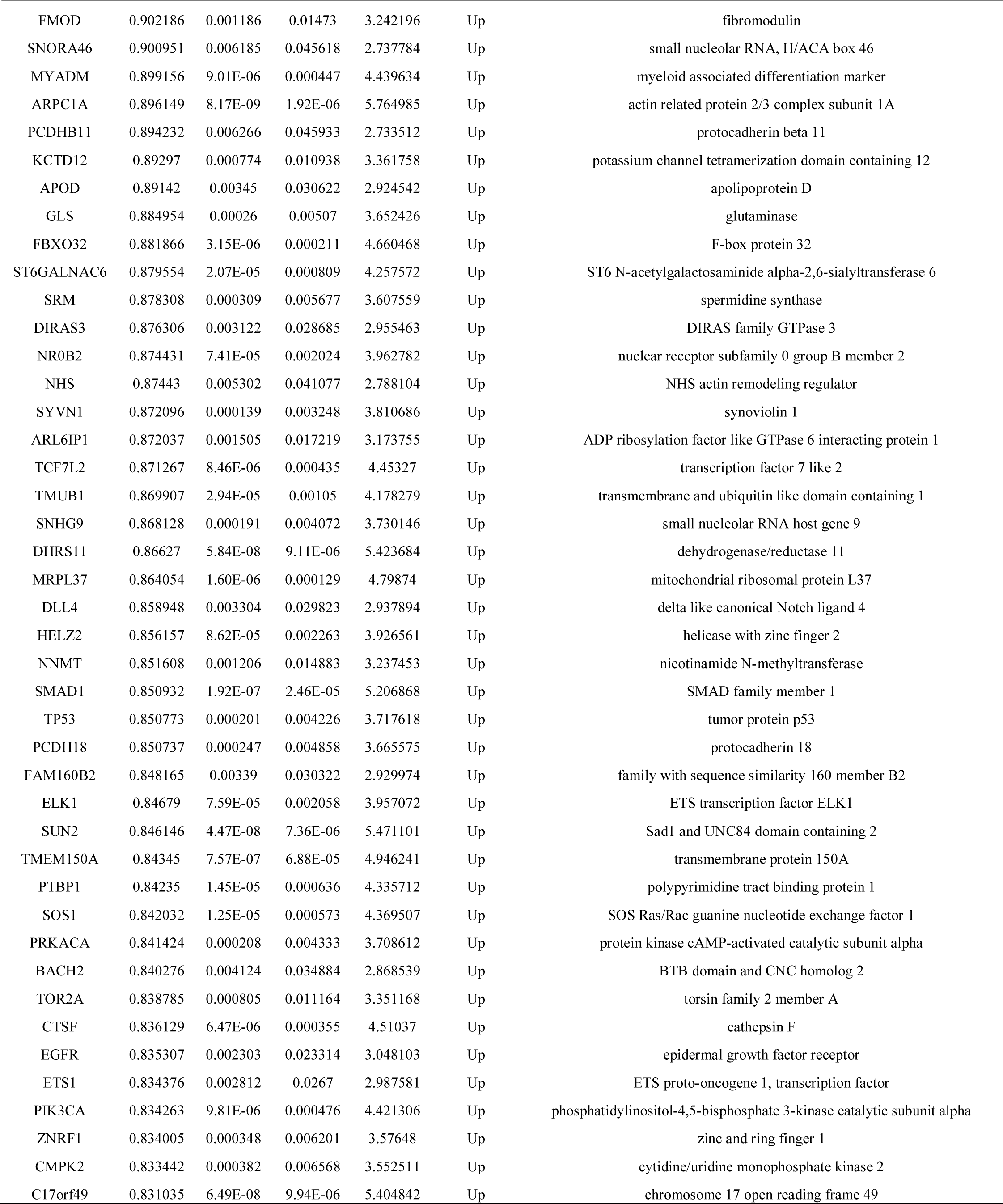
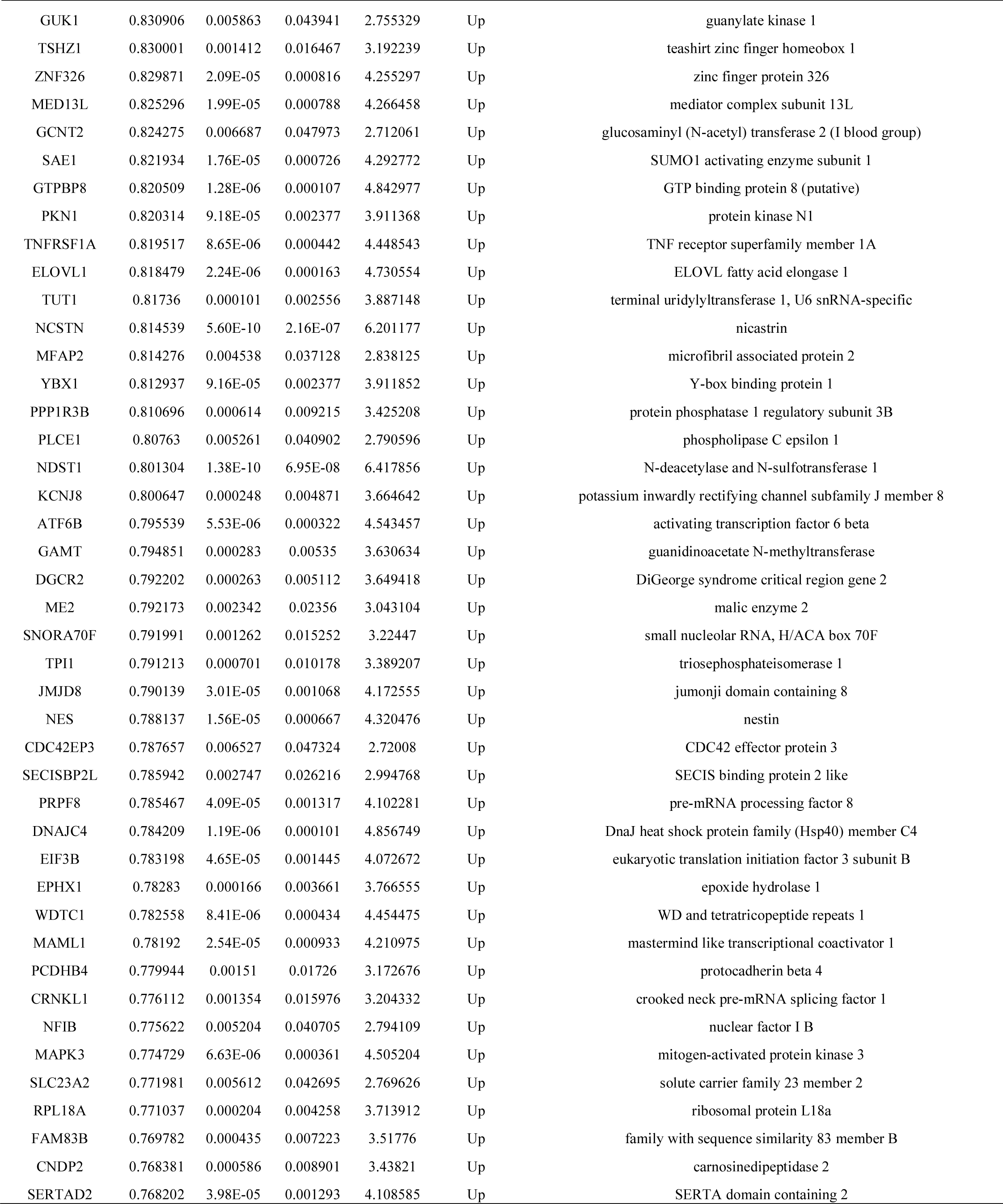
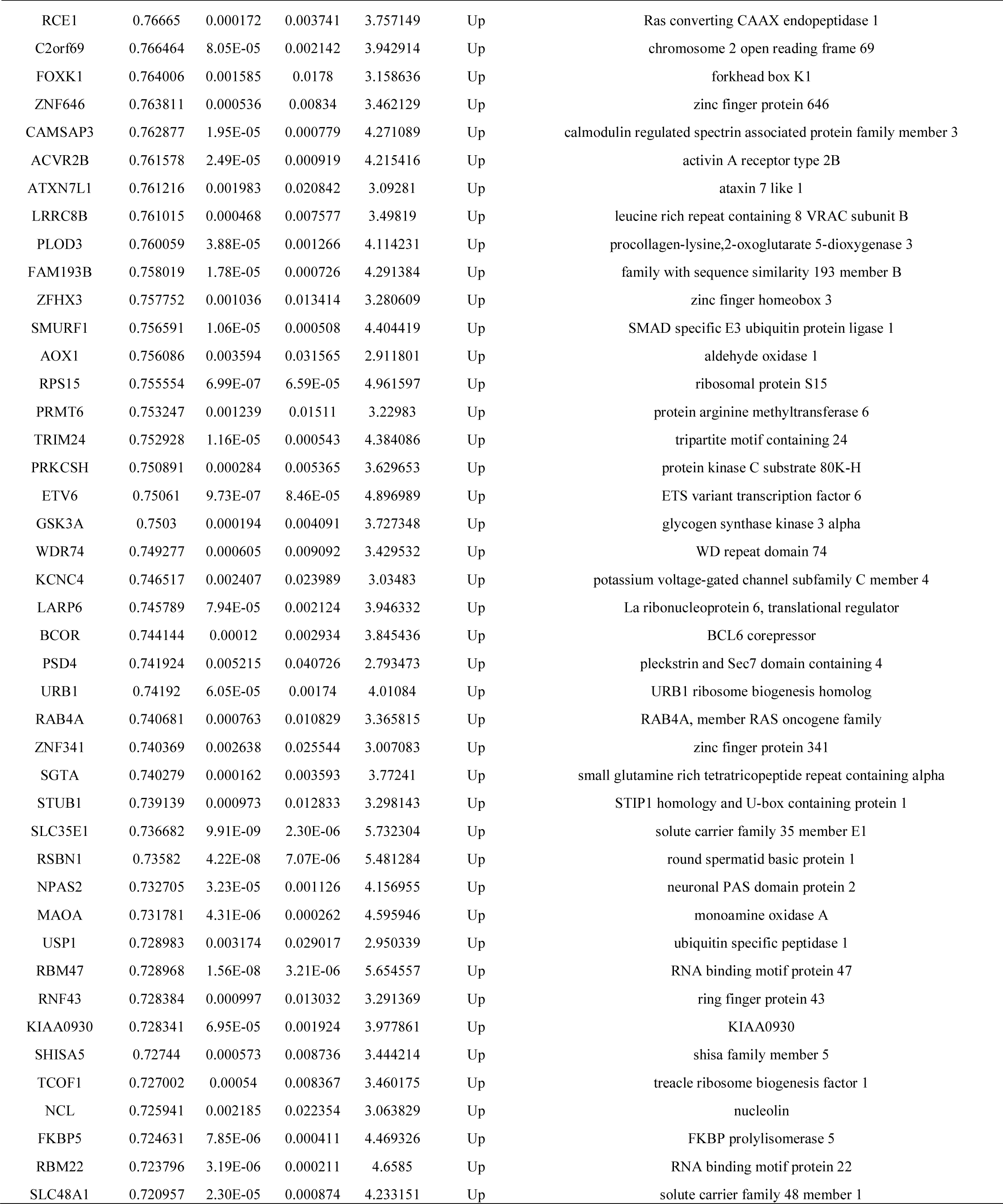
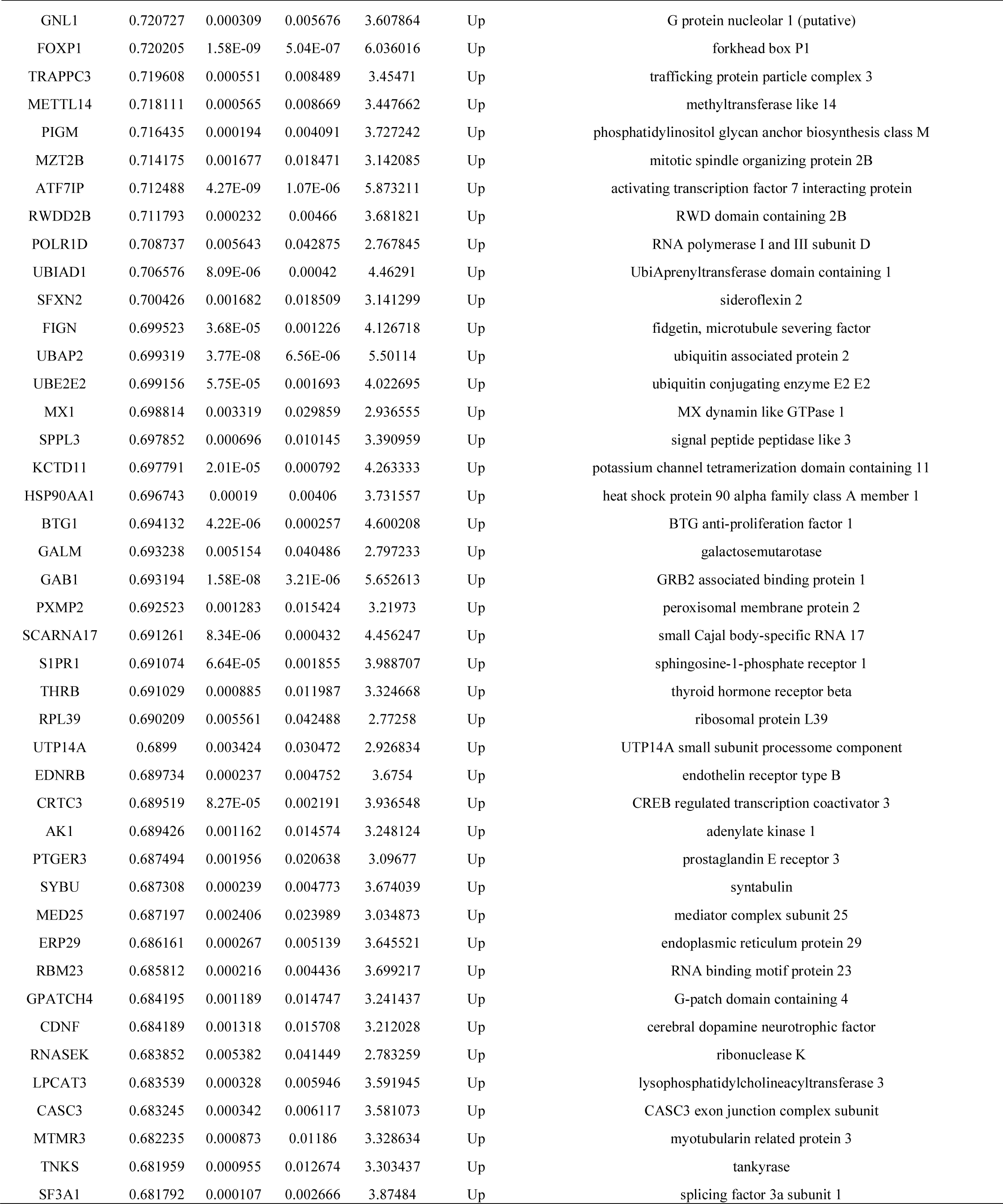
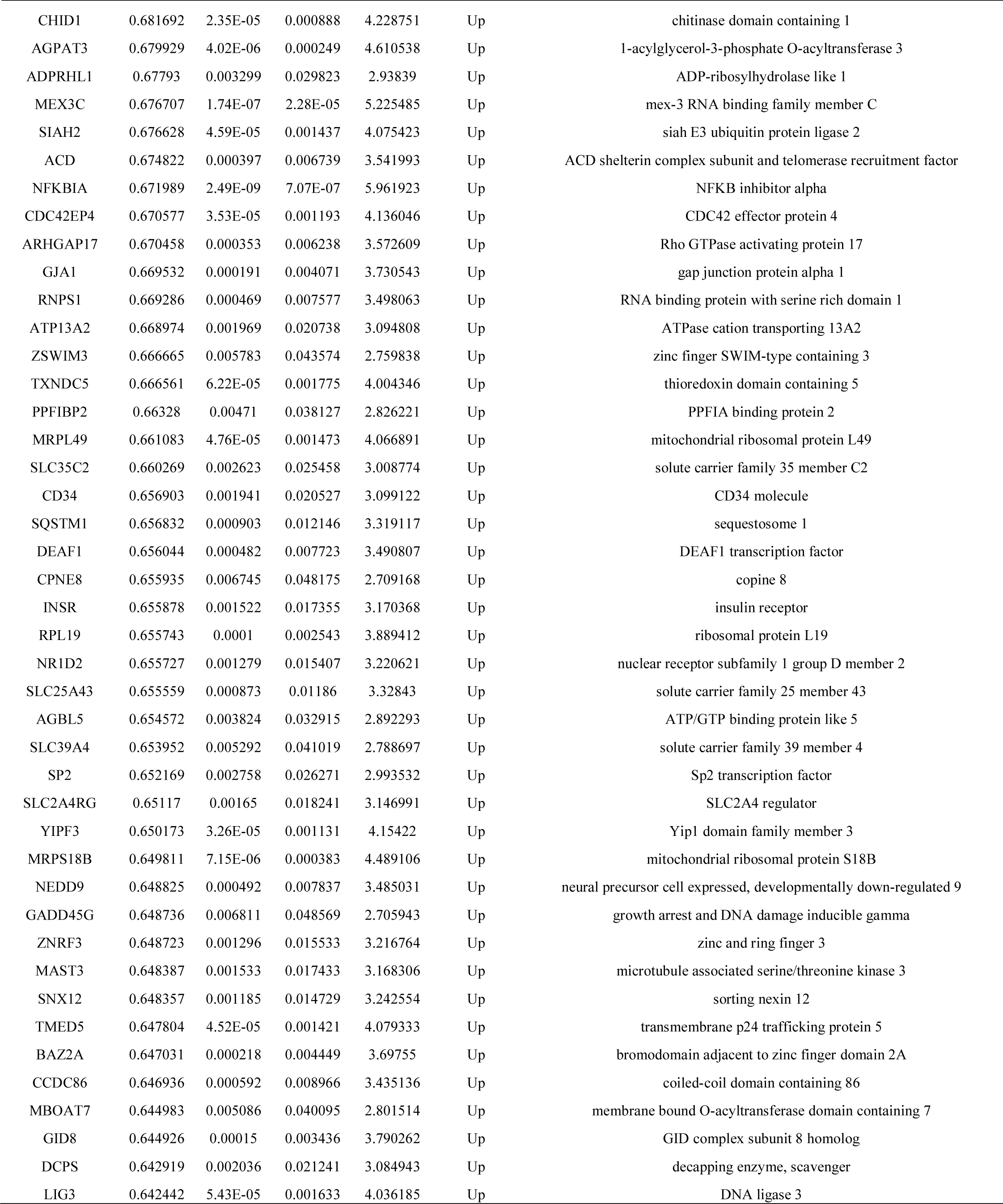
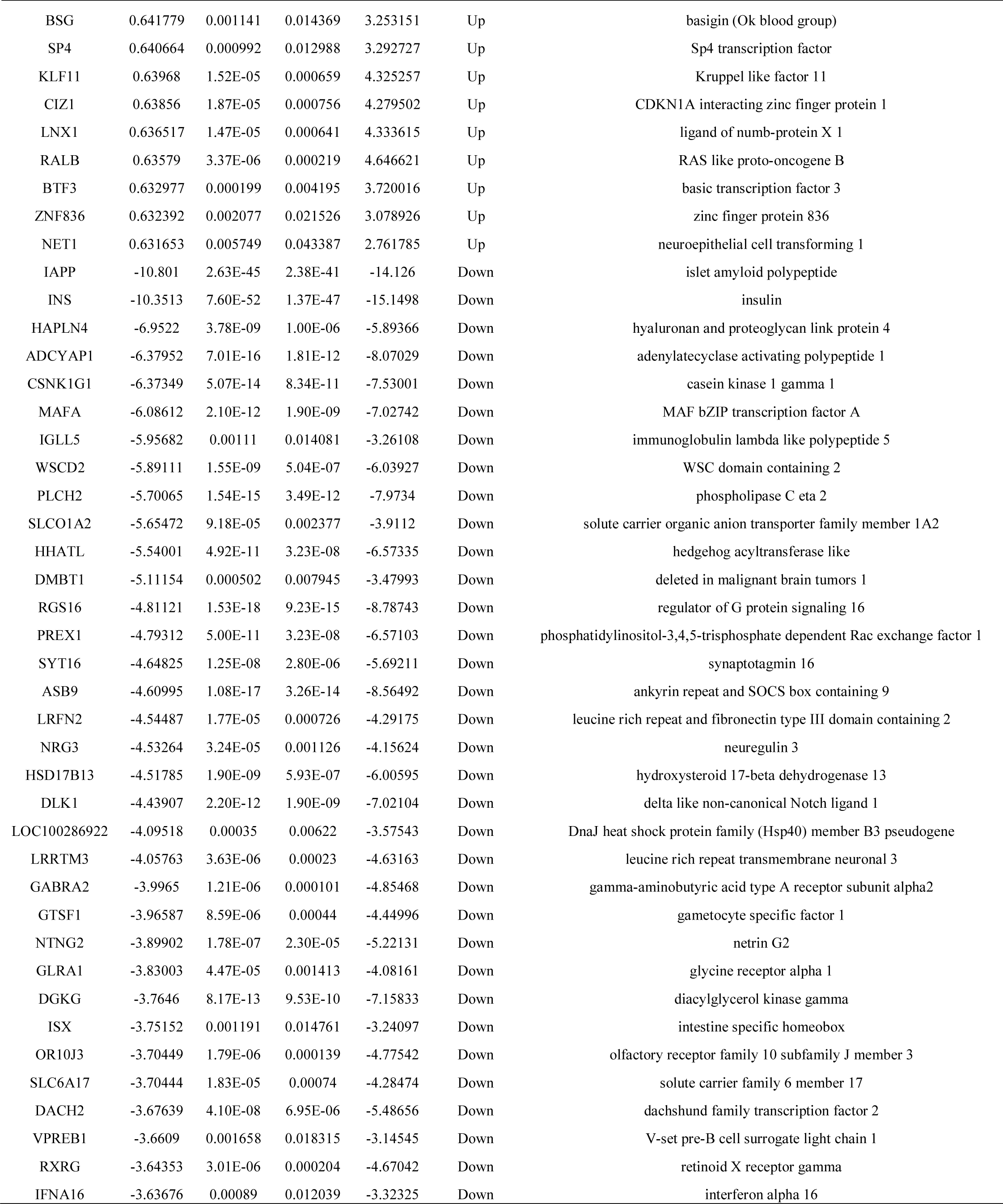
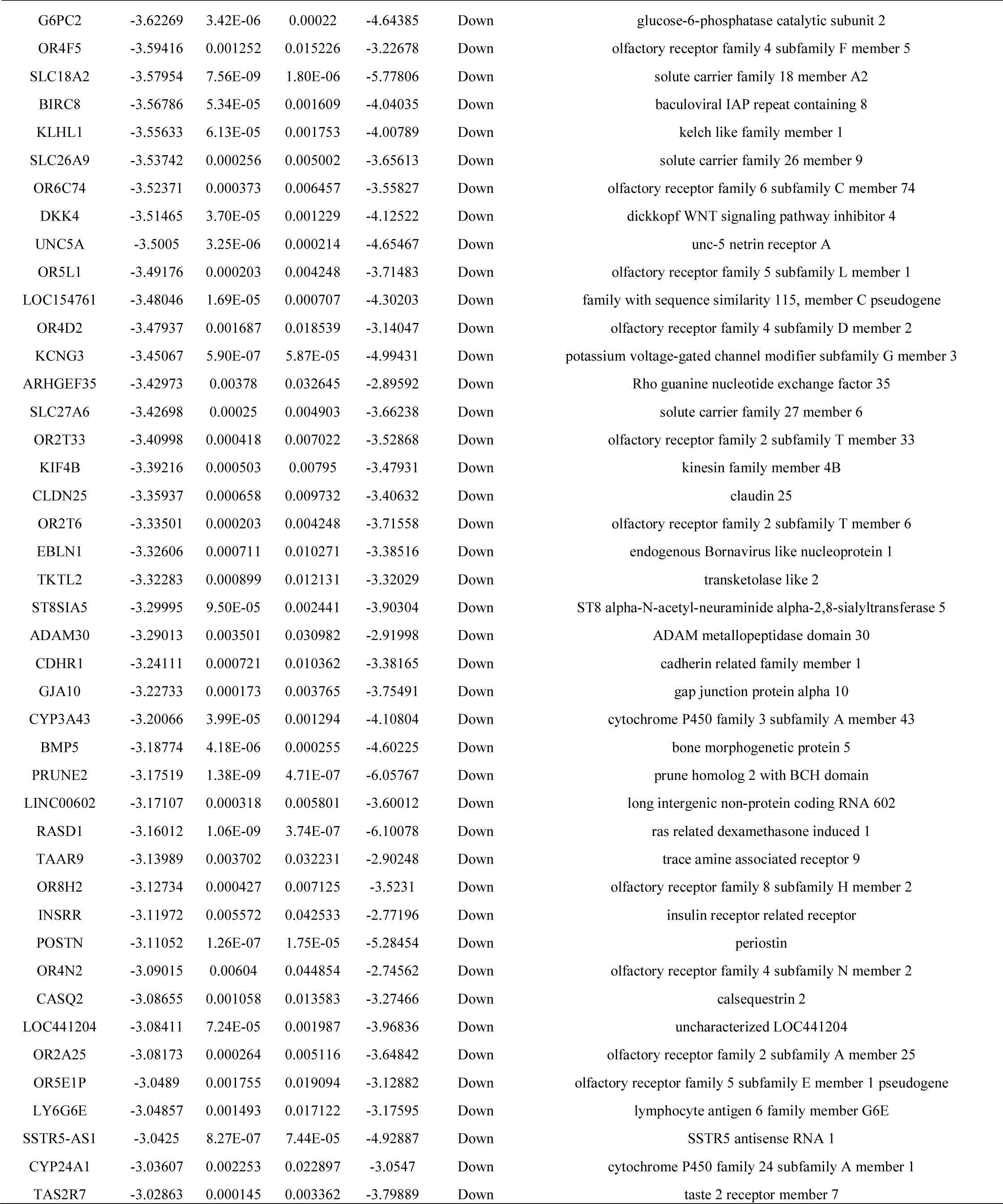
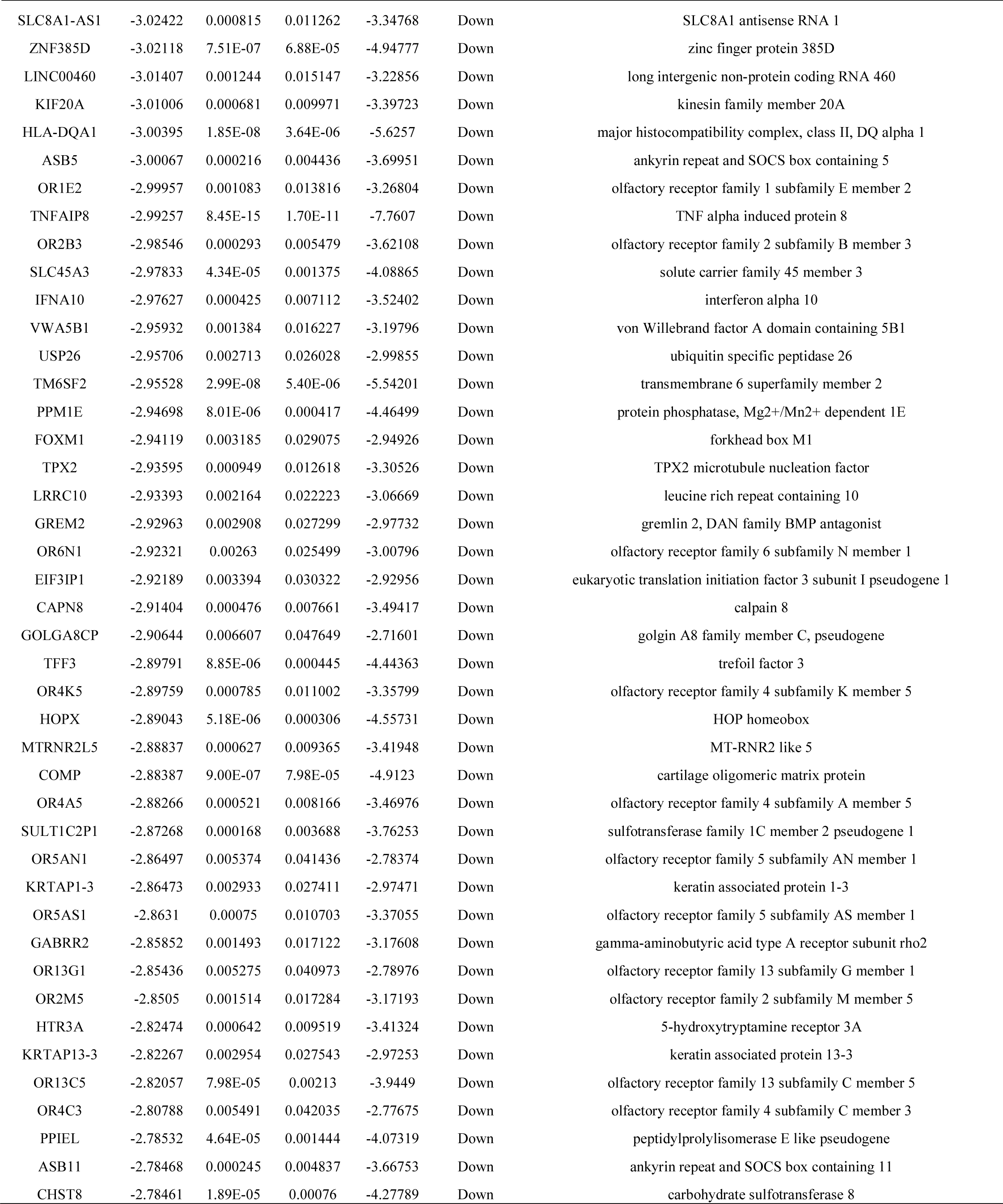
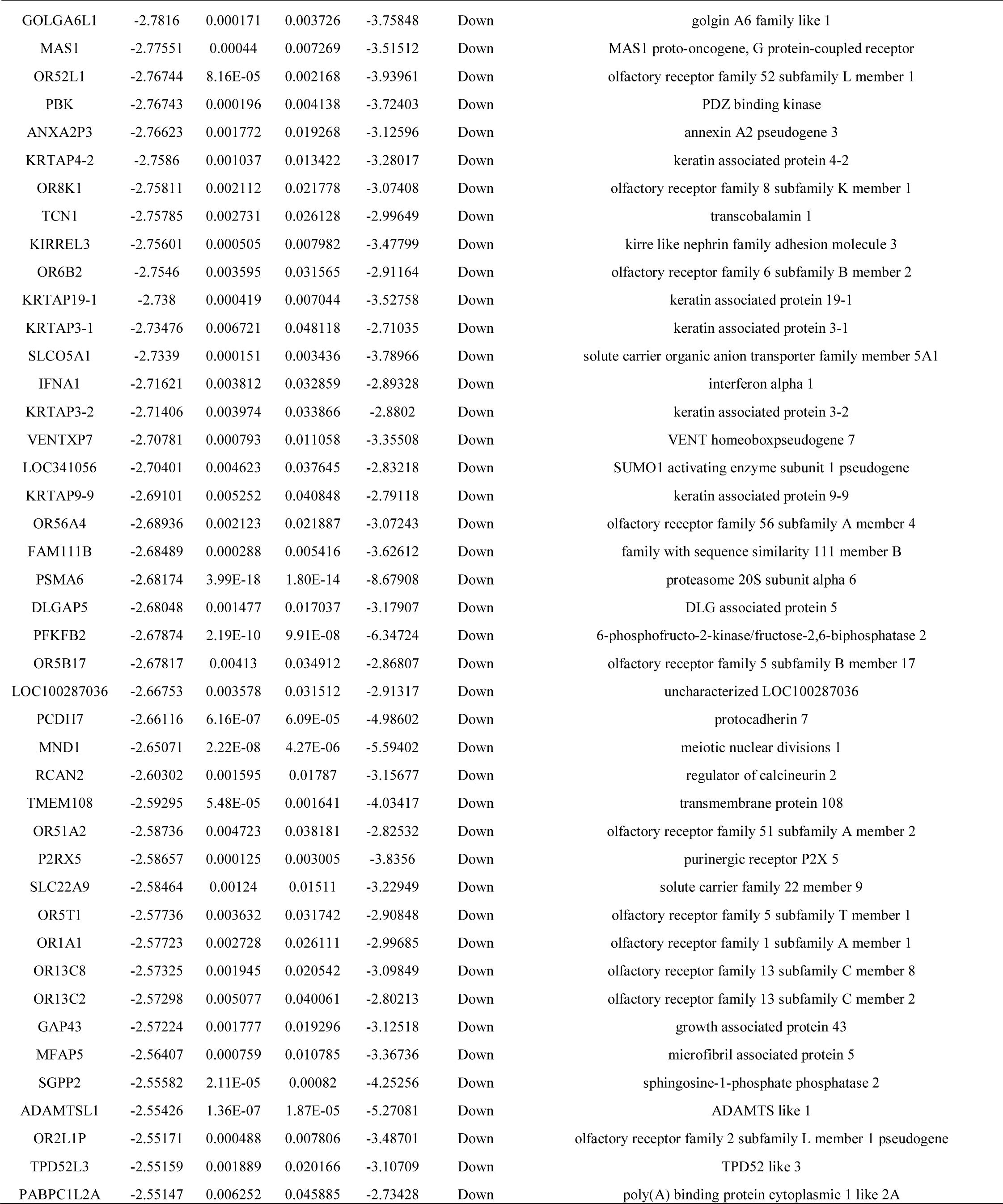
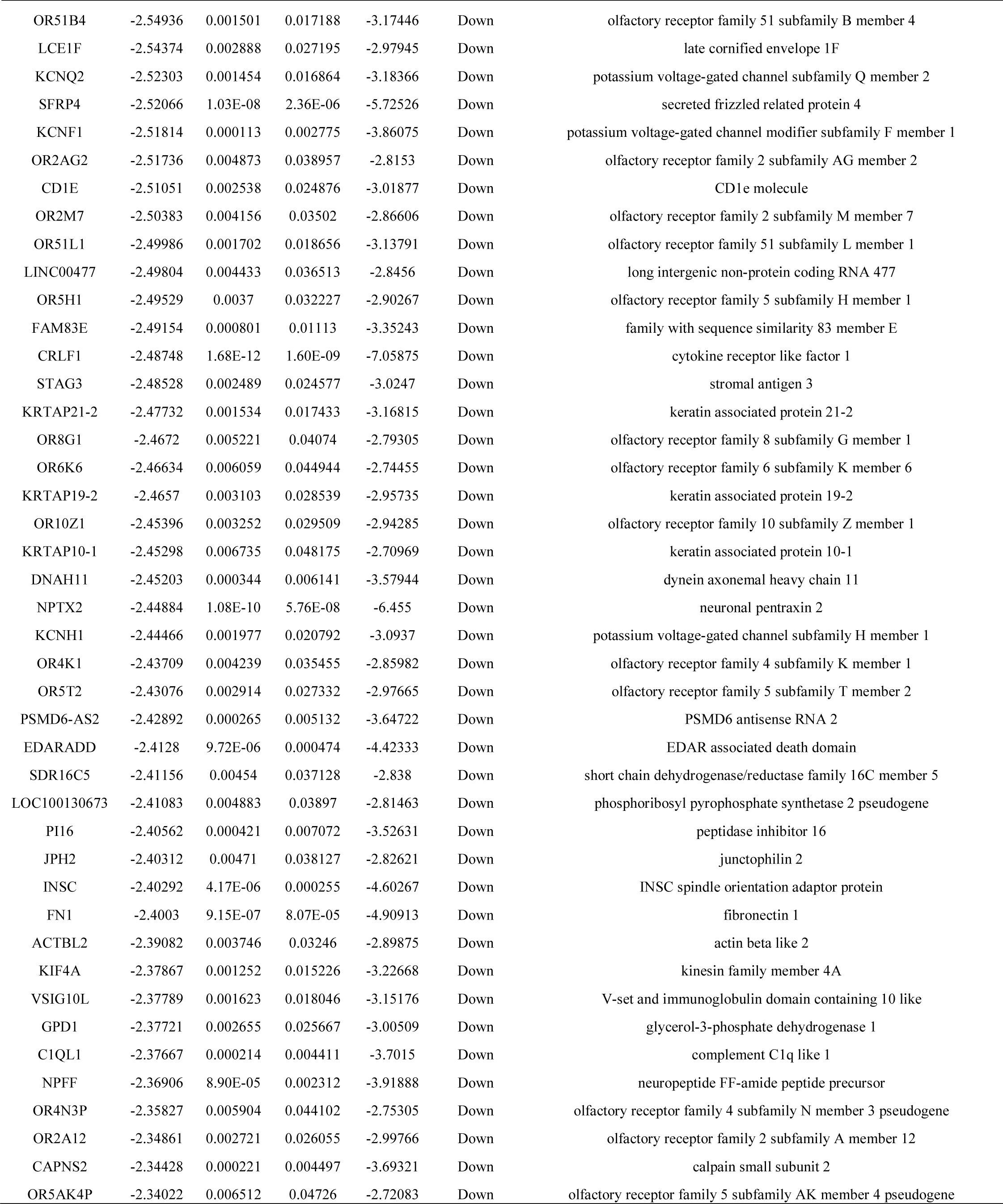
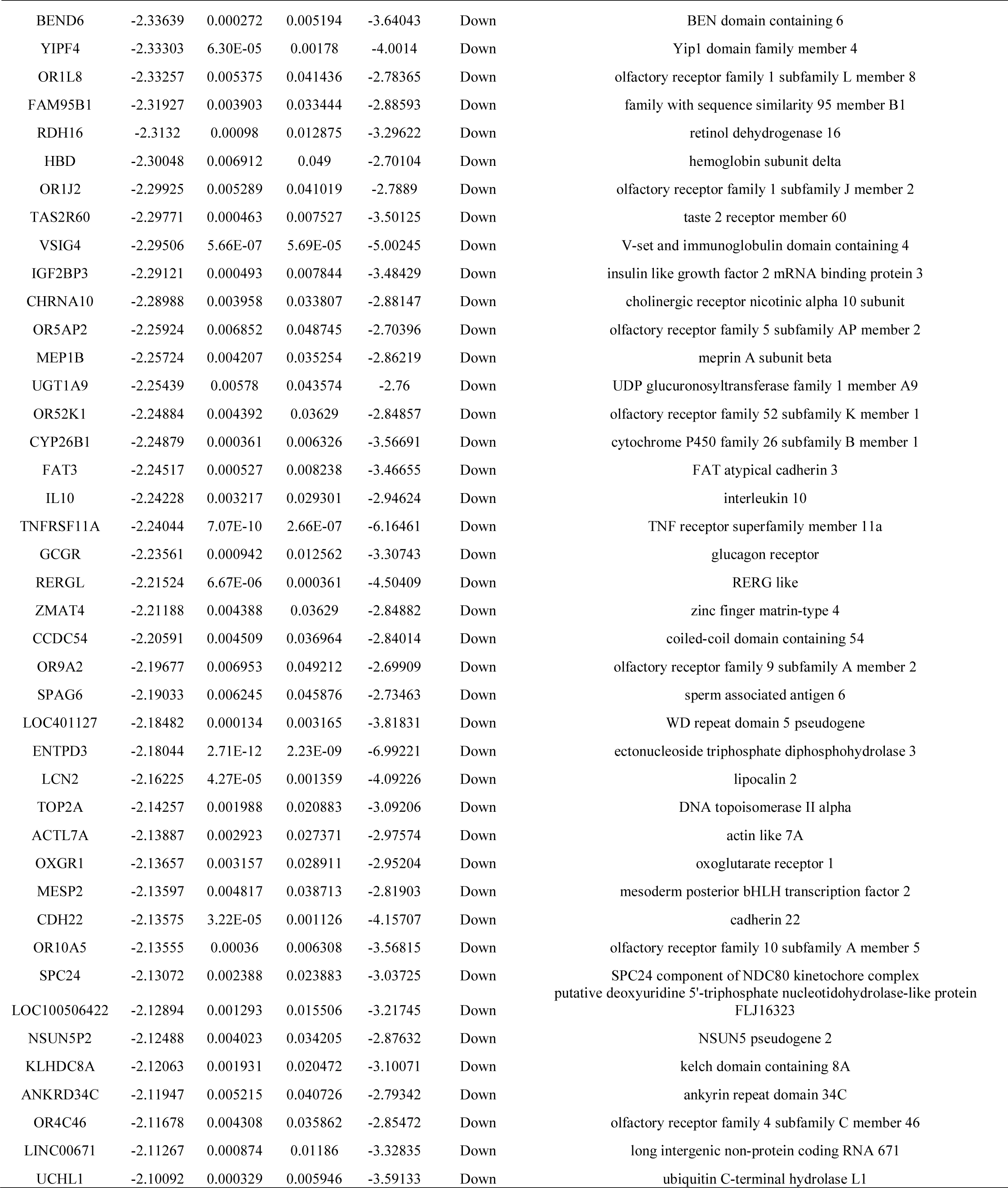
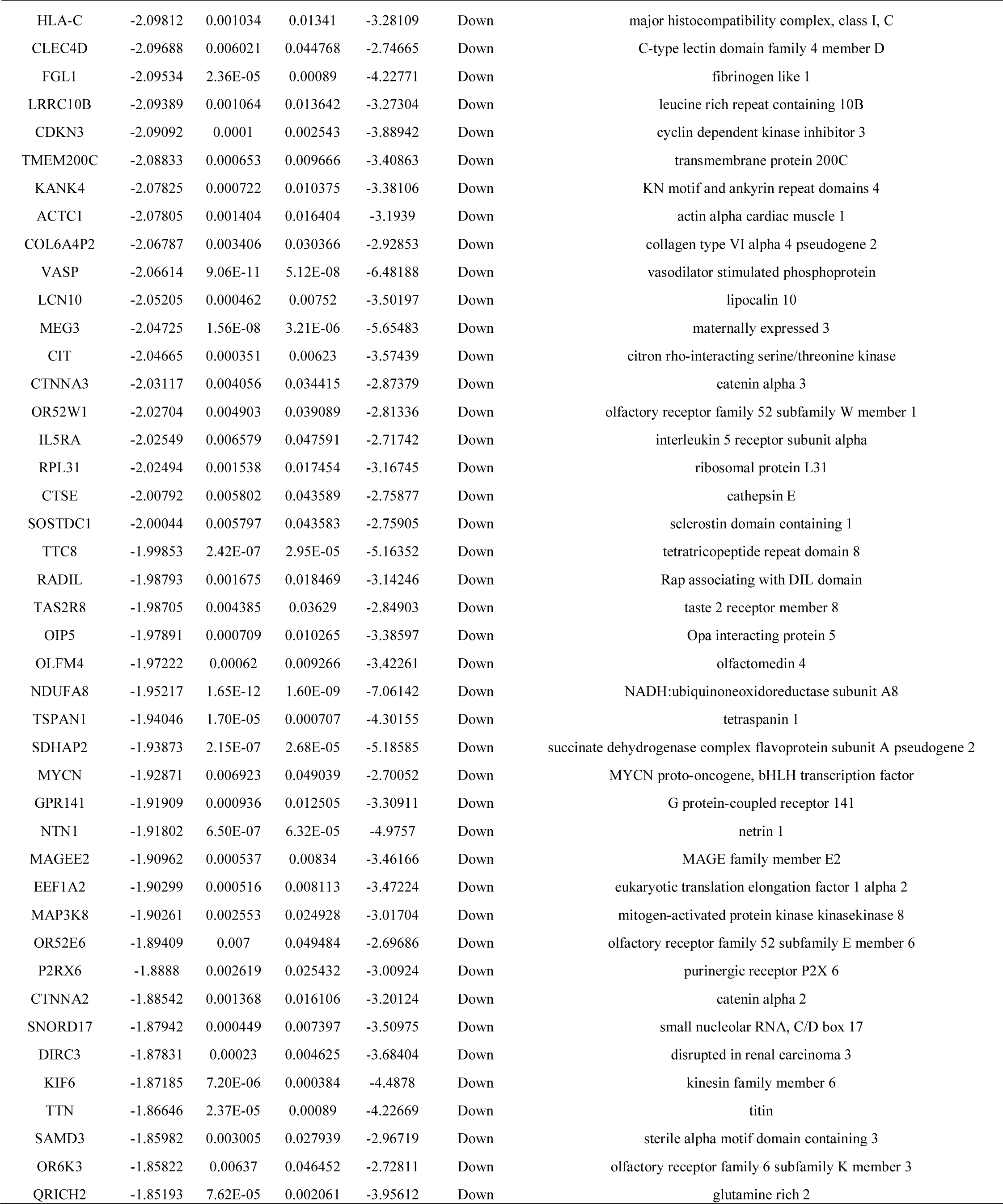
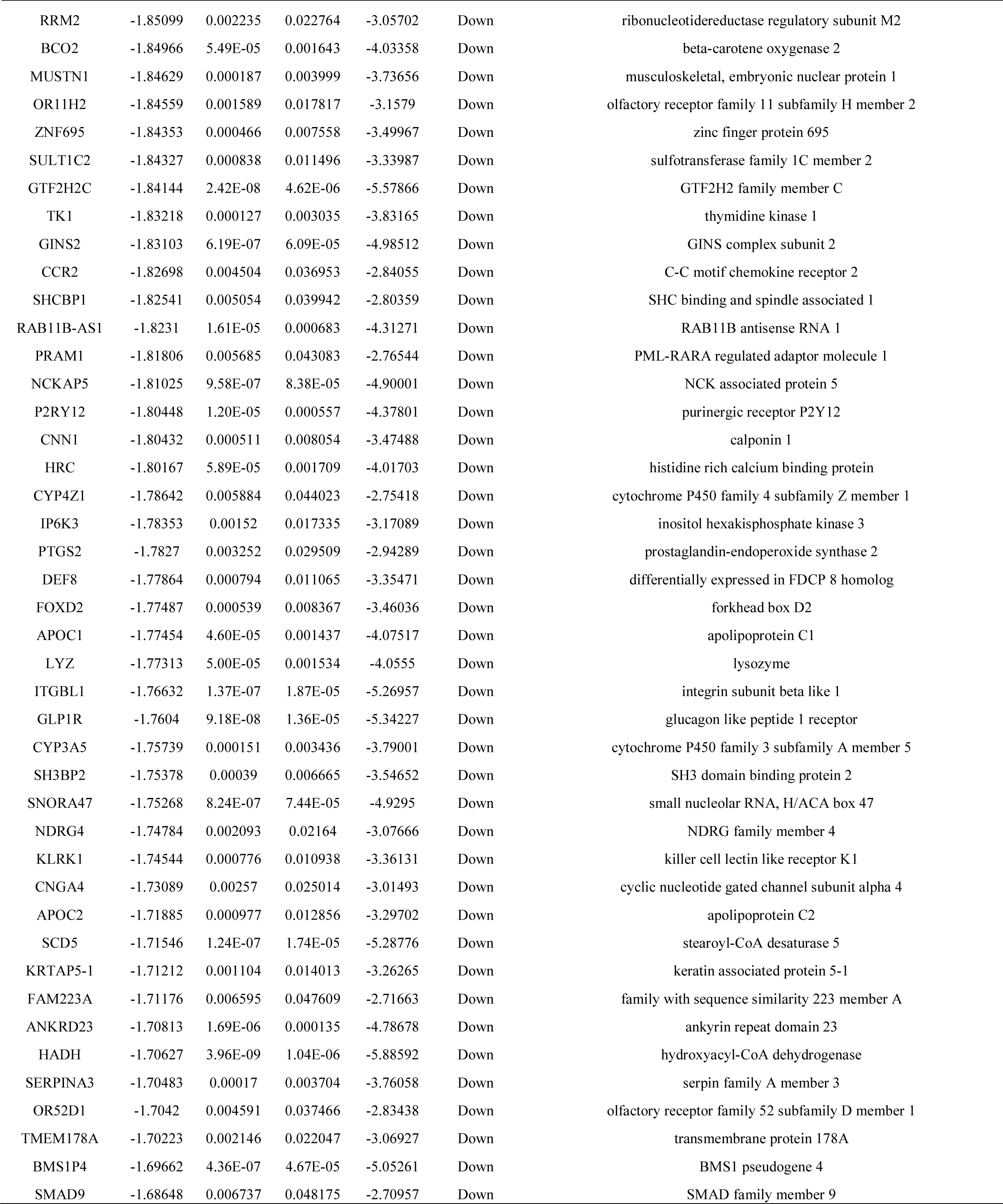
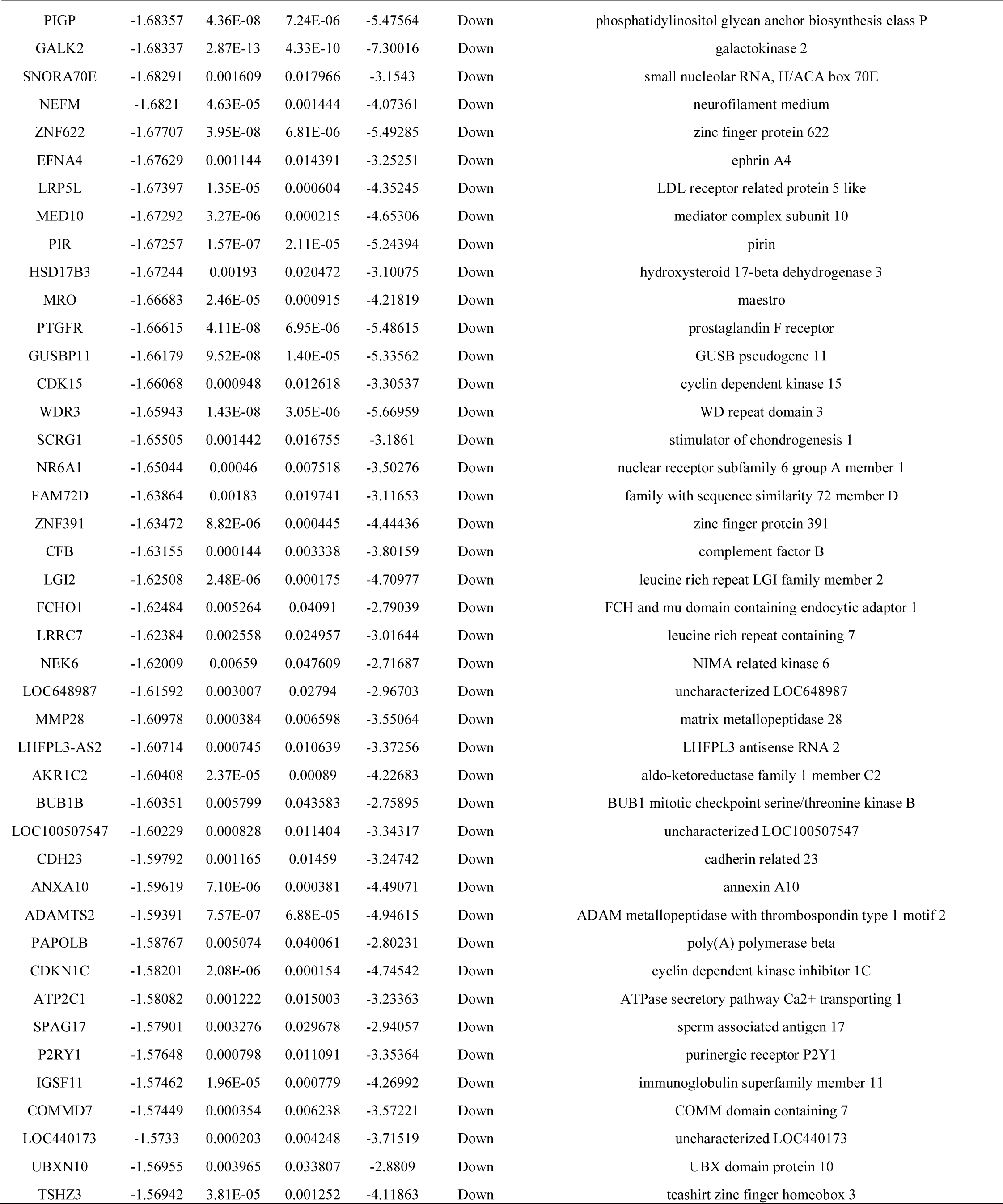
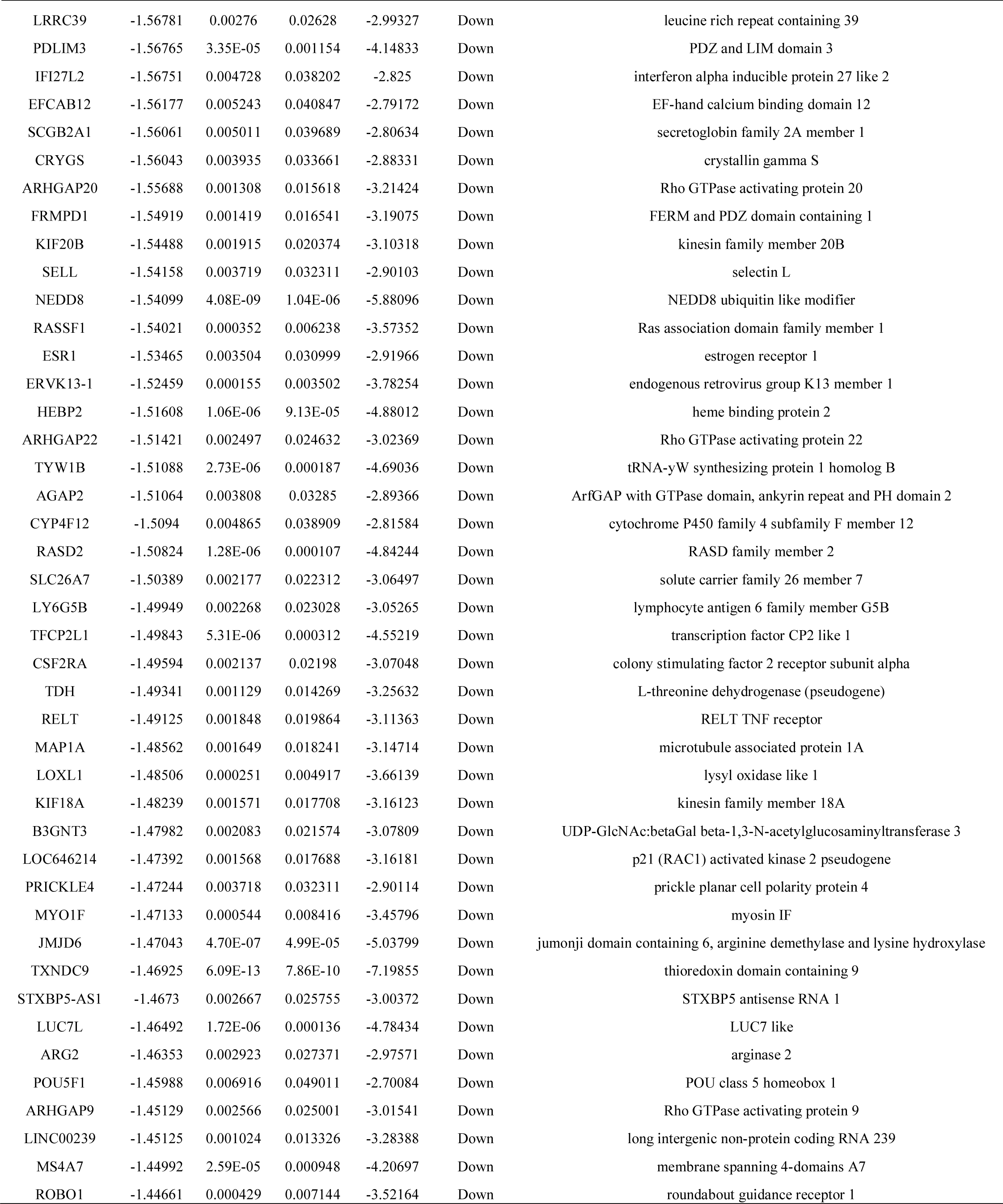
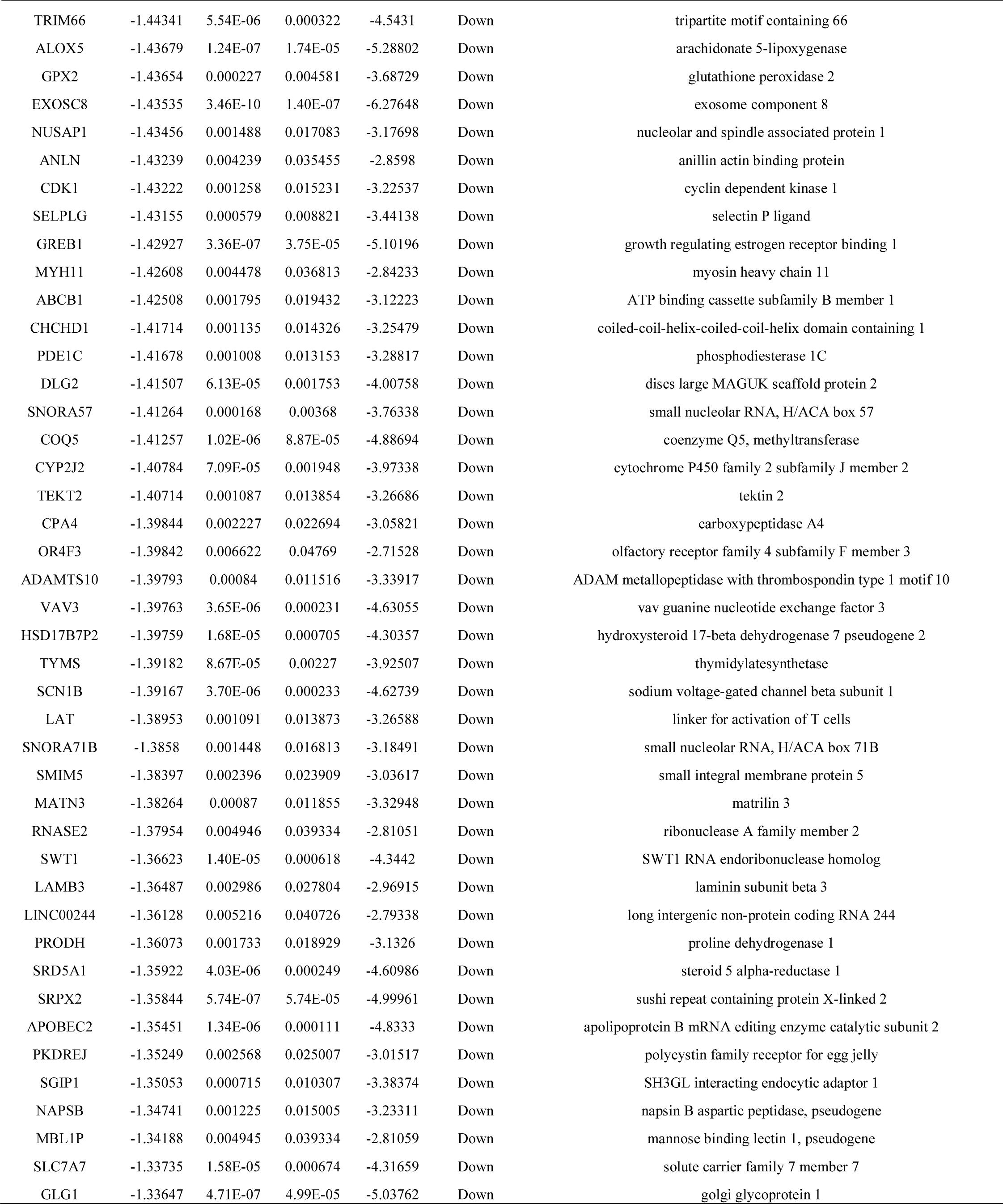
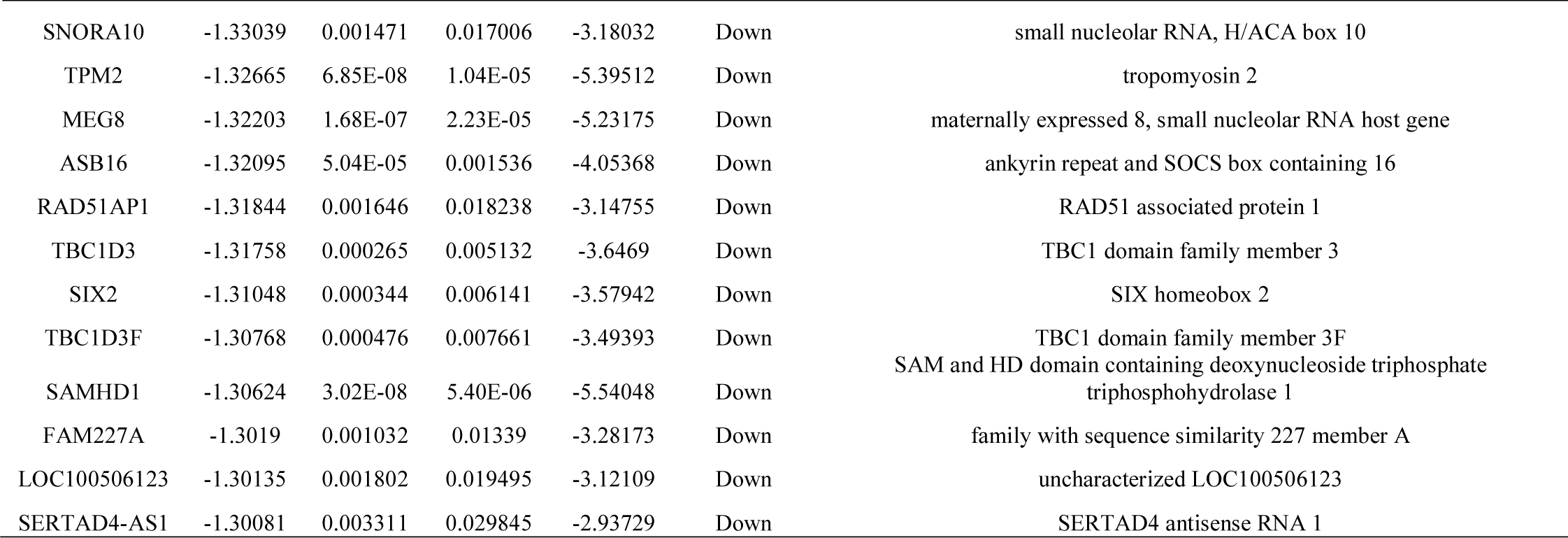
The statistical metrics for key differentially expressed genes (DEGs)

### GO and REACTOME pathway enrichment analysis of DEGs

To characterize the functional roles of the above DEGs, we used GO (Table 3) and REACTOME pathway (Table 4) enrichment analyses. The BP category of the GO analysis results showed that up regulated genes were significantly enriched in multicellular organism development and nitrogen compound metabolic process. For CC, these up regulated were enriched in membrane-enclosed lumen and nuclear lumen. Moreover, up regulated genes were significantly enriched in protein binding and transcription regulator activity in the MF categories. In addition, the most significantly enriched GO terms for down regulated genes were detection of stimulus and multicellular organismal process (BP), cell periphery and plasma membrane (CC), and transmembrane signaling receptor activity and molecular transducer activity (MF). According to REACTOME pathway enrichment analysis, up regulated genes were significantly enriched in diseases of signal transduction by growth factor receptors and second messengers and formation of the cornified envelope. Down regulated genes were enriched in olfactory signaling pathway and sensory perception.

**Table 3.**
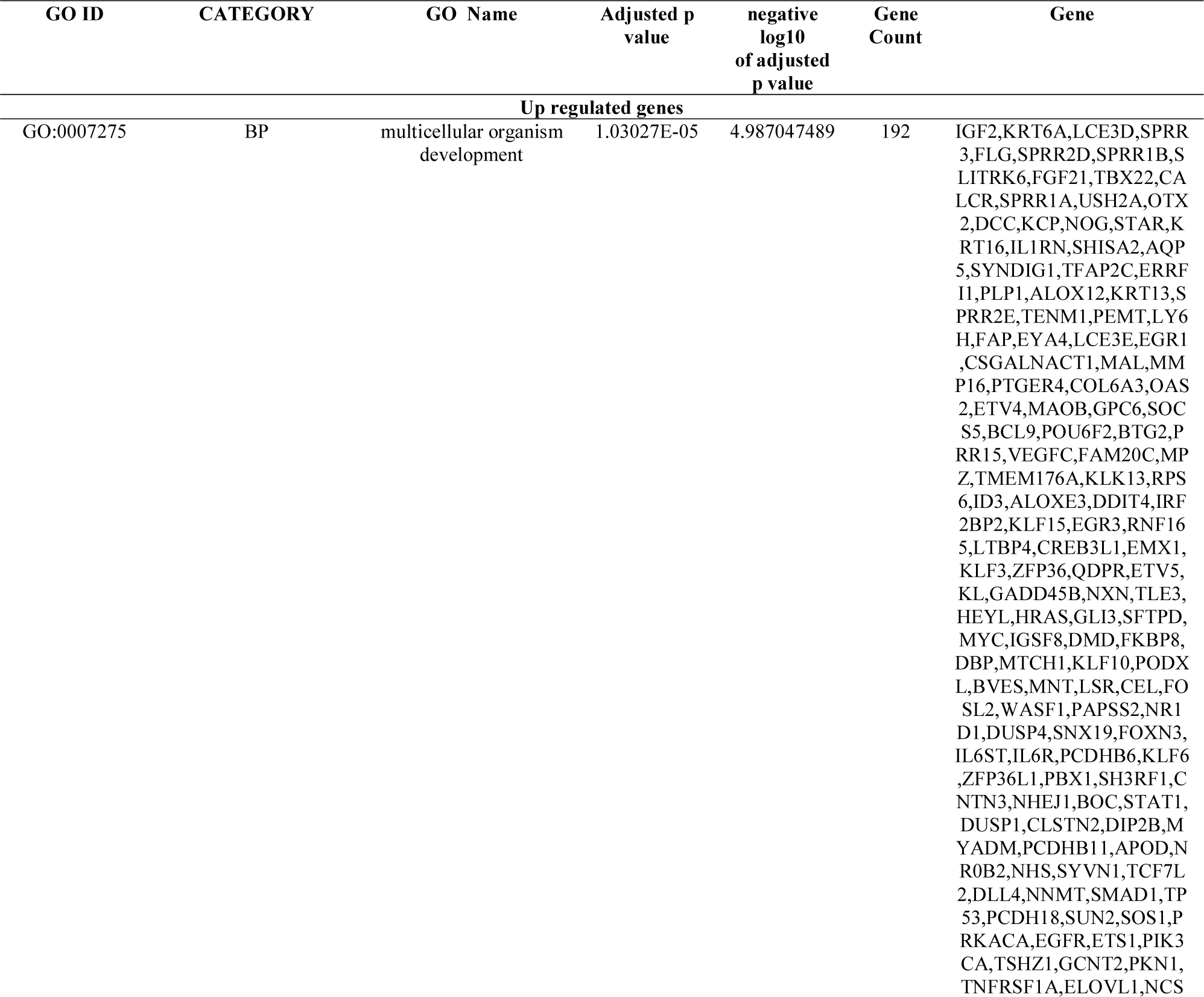
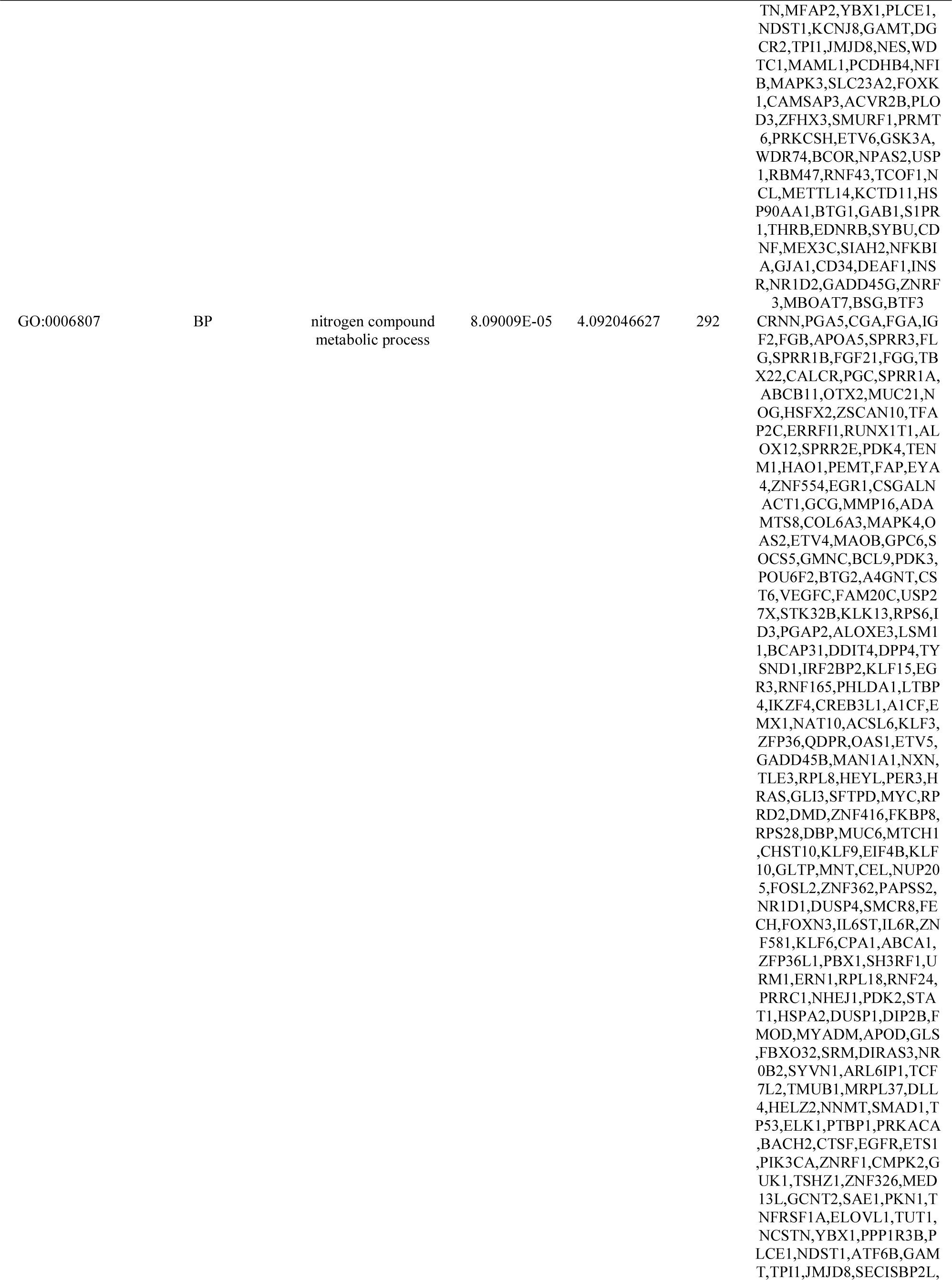
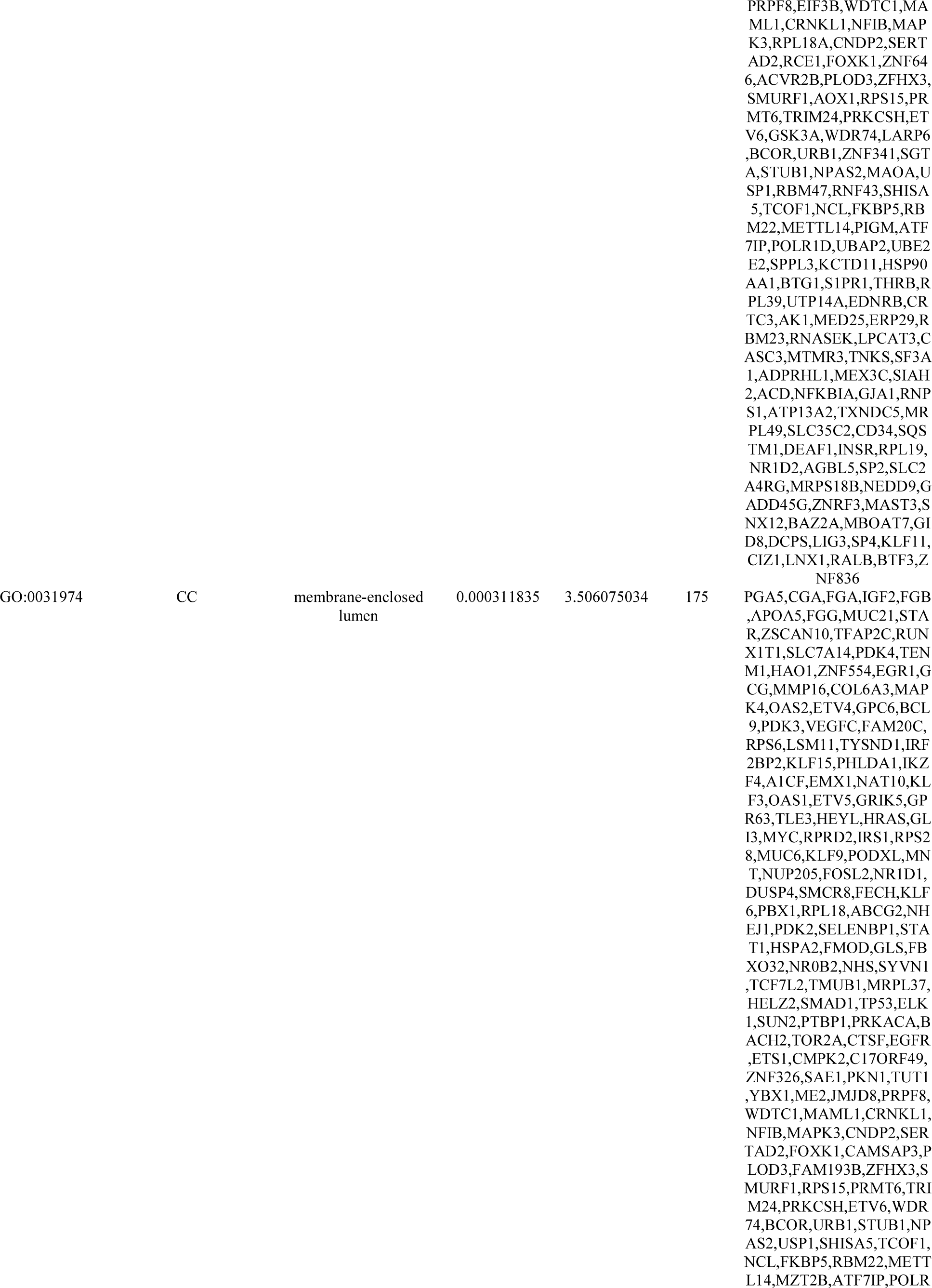
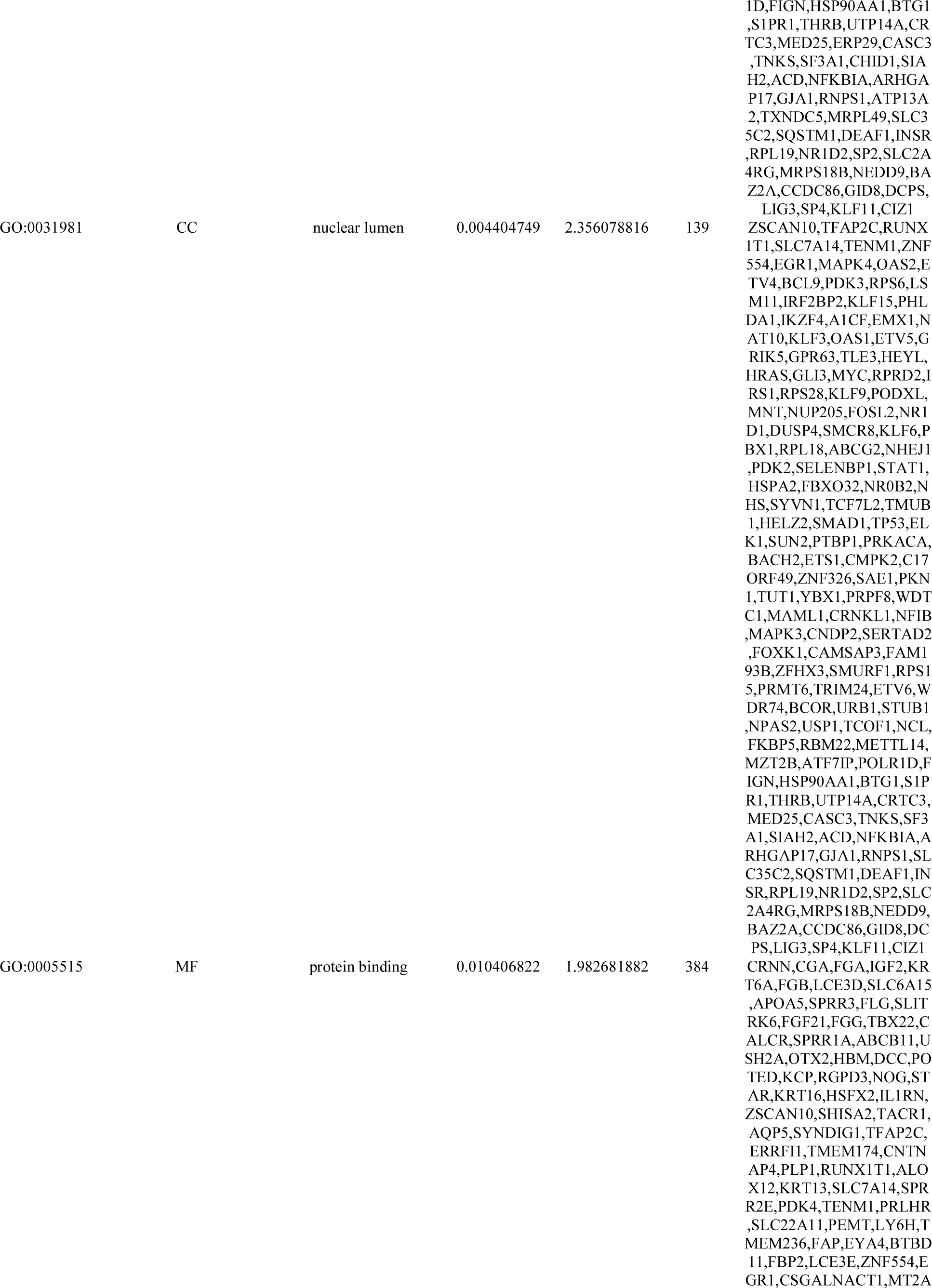
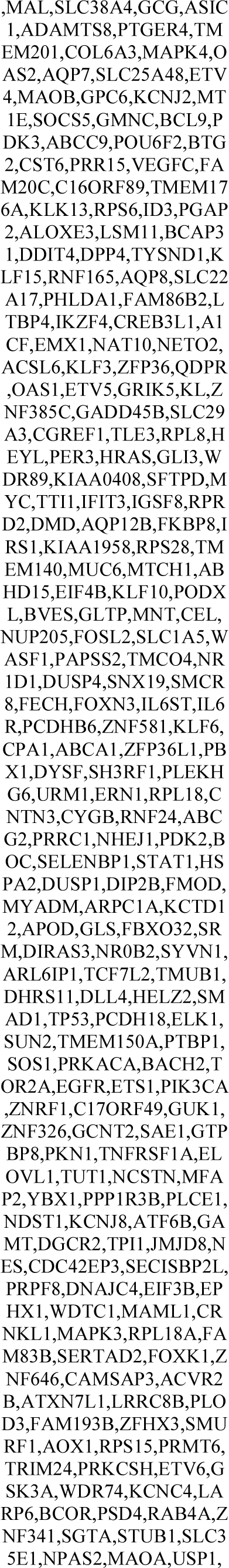
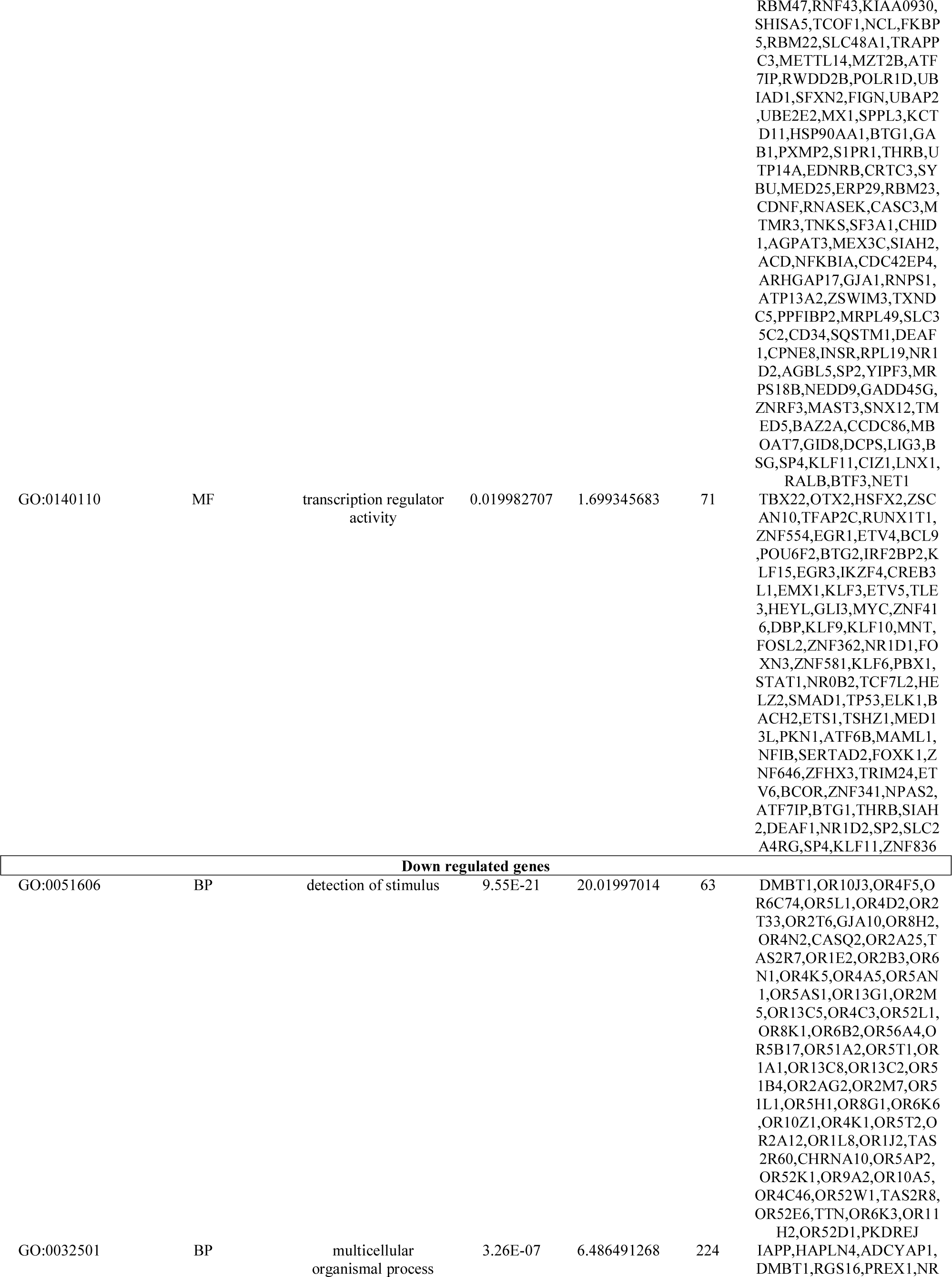
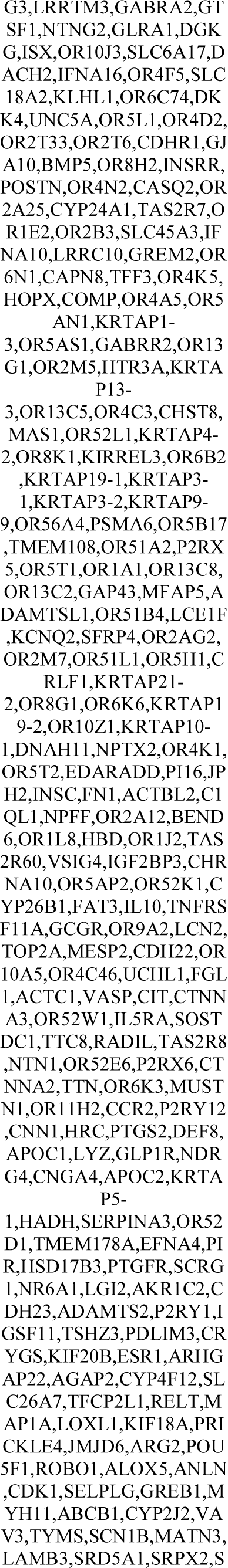
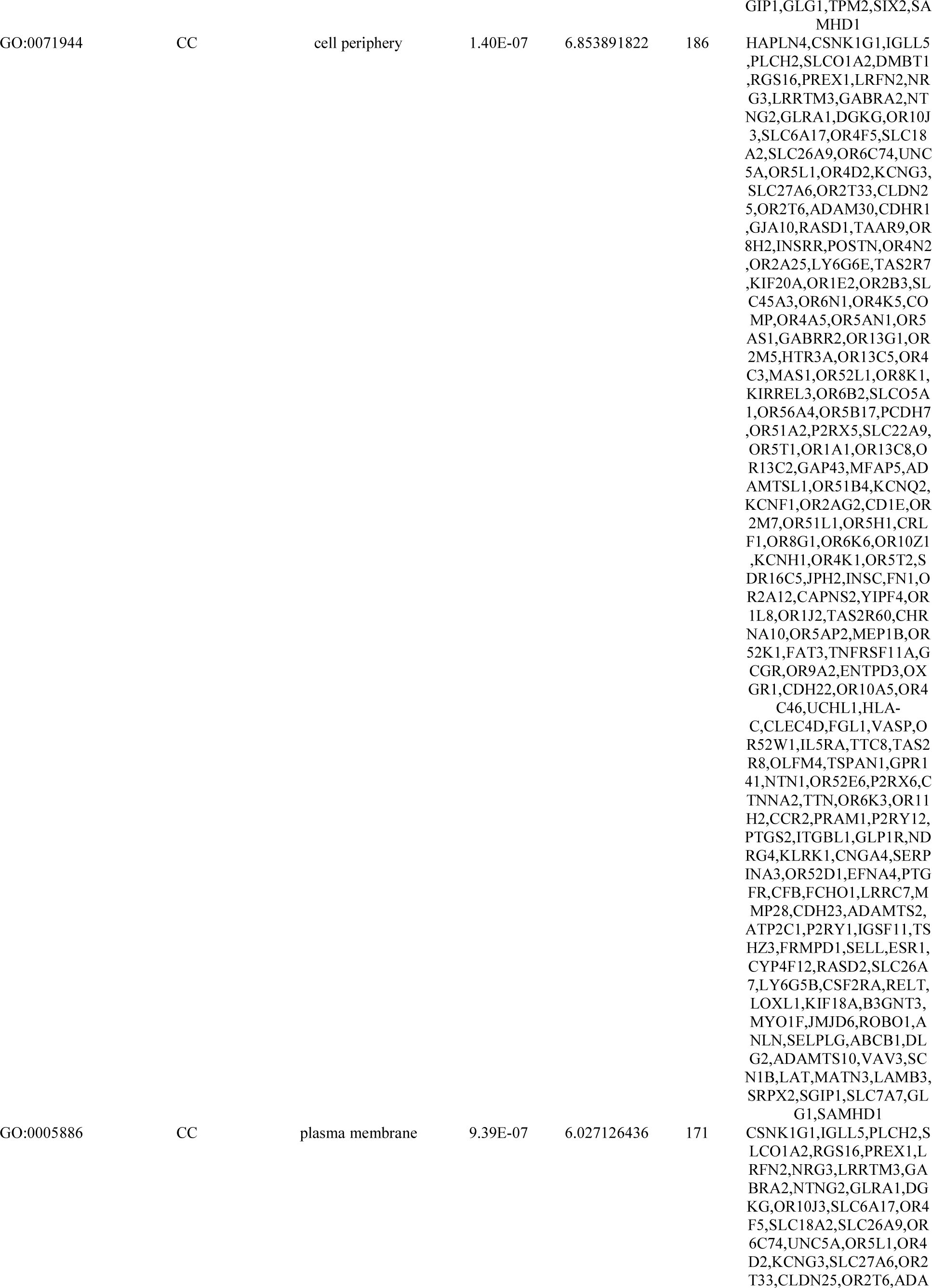
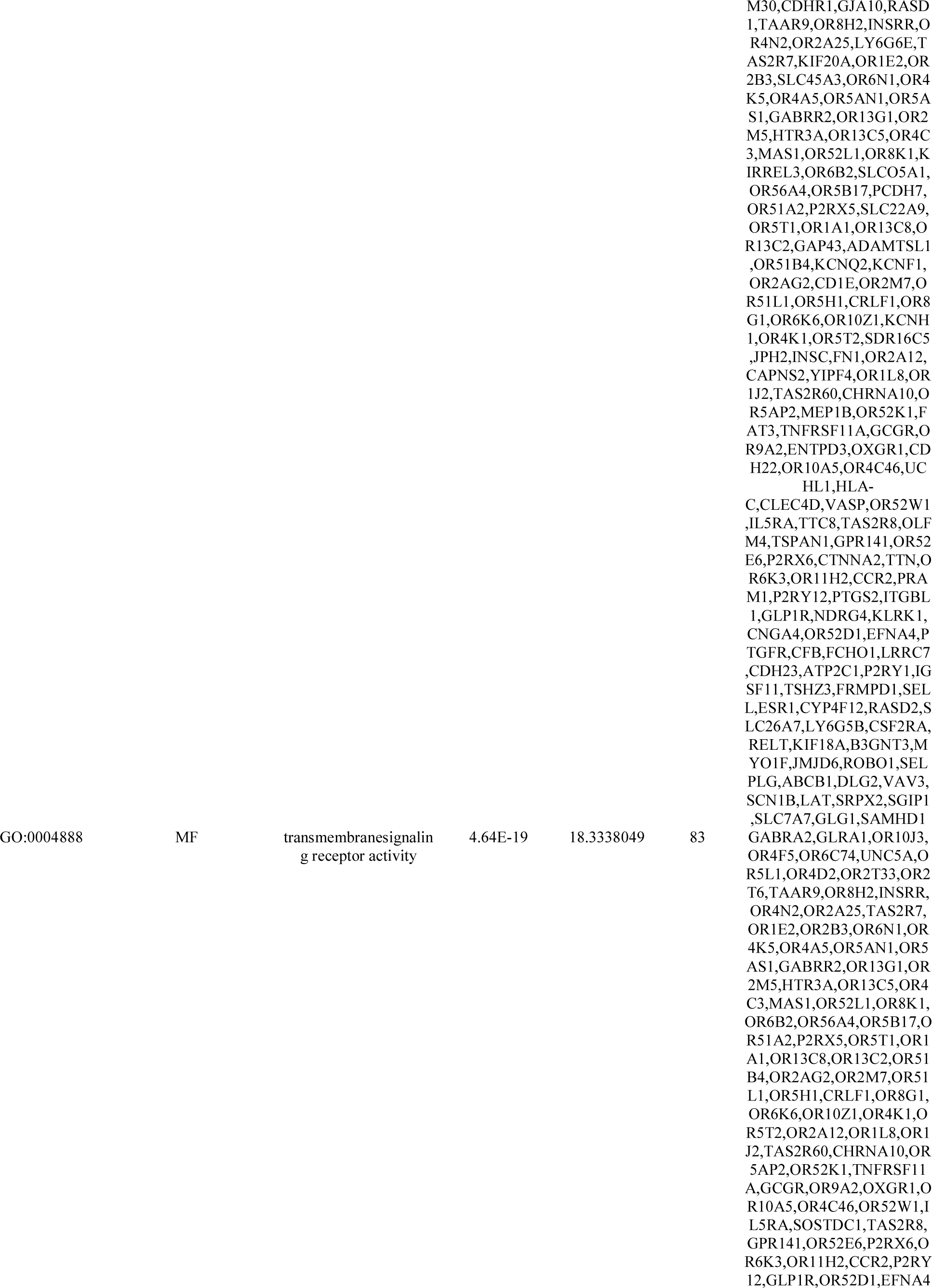
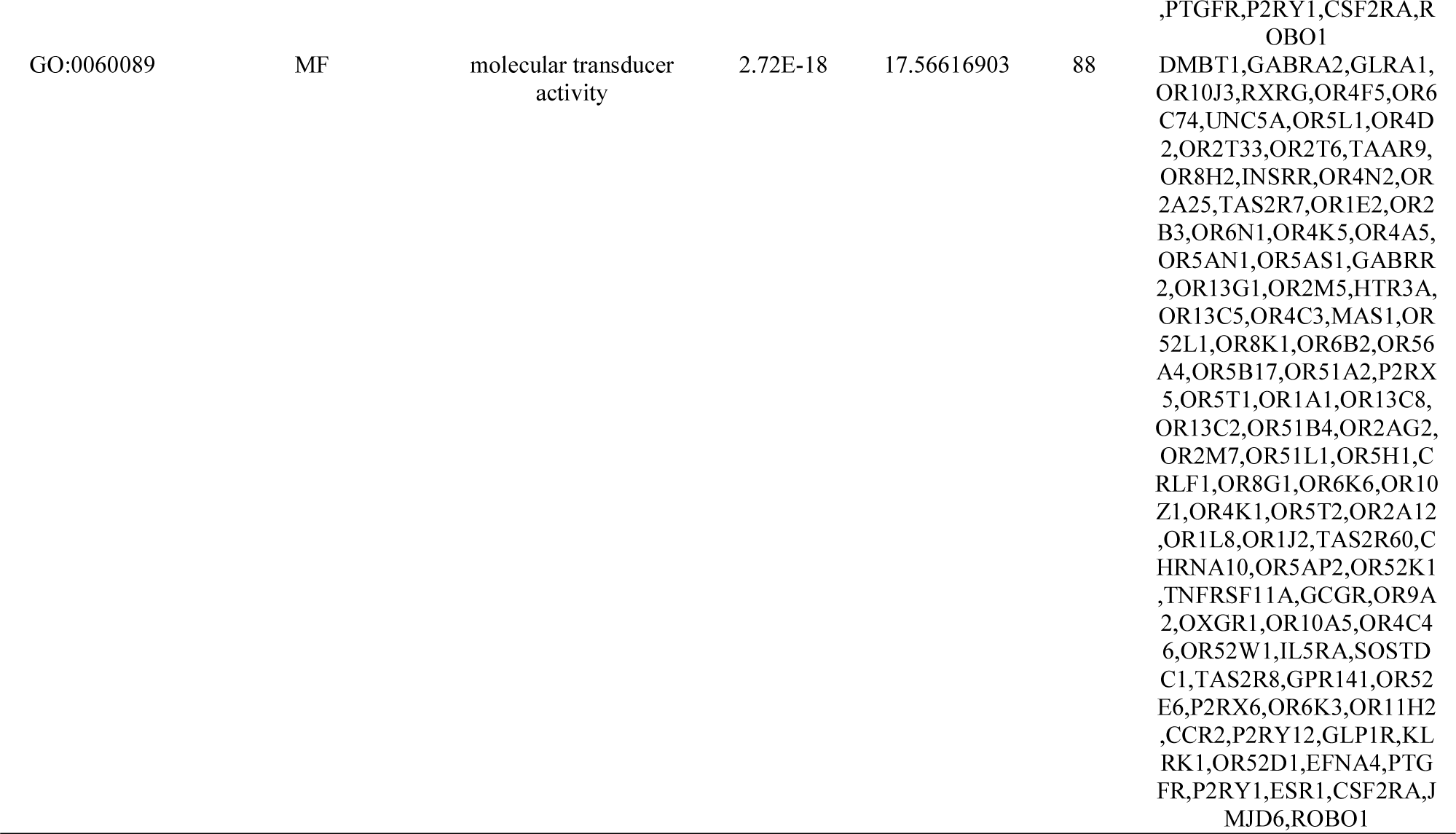
The enriched GO terms of the up and down regulated differentially expressed genes

**Table 4.**
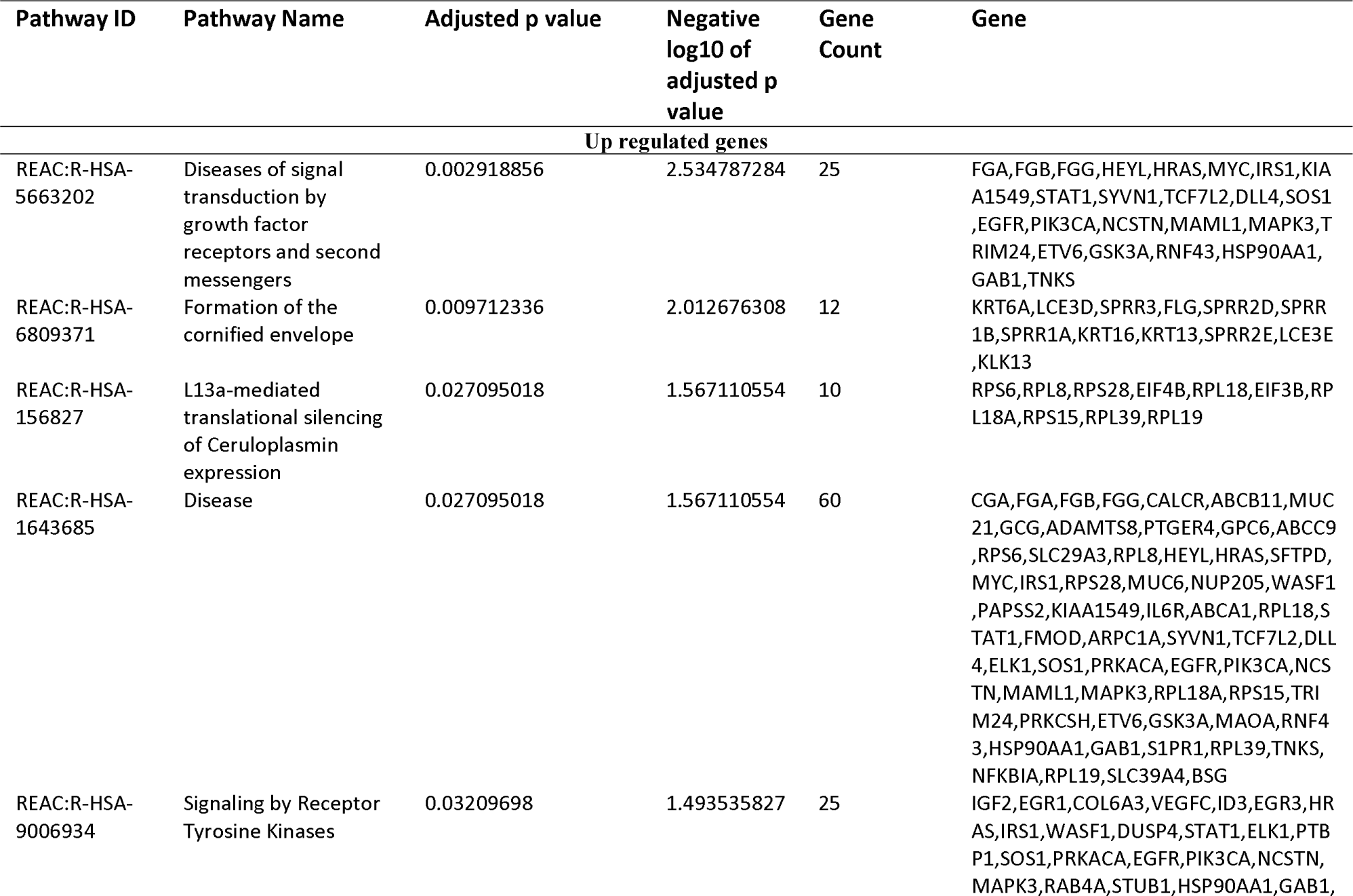
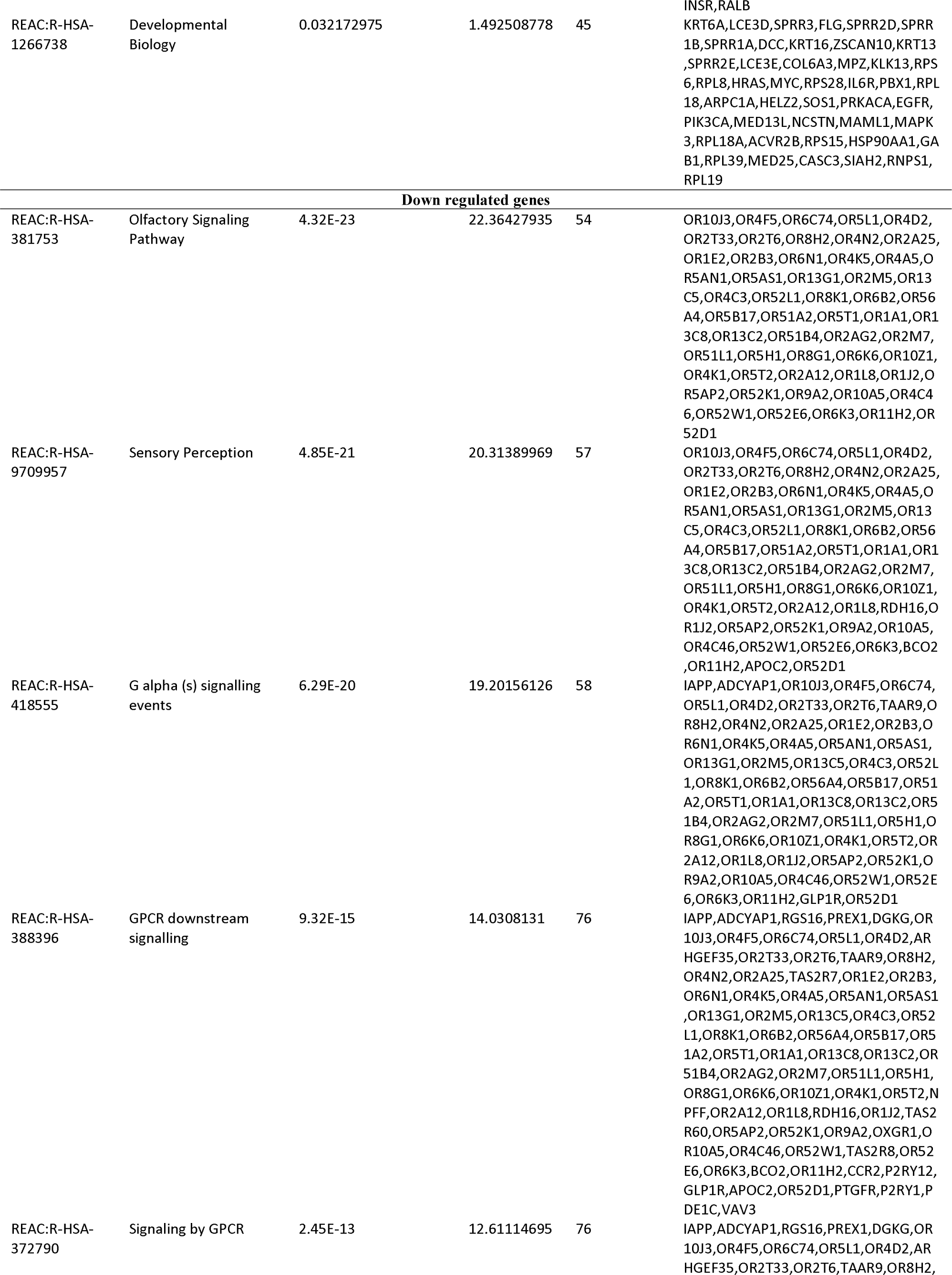
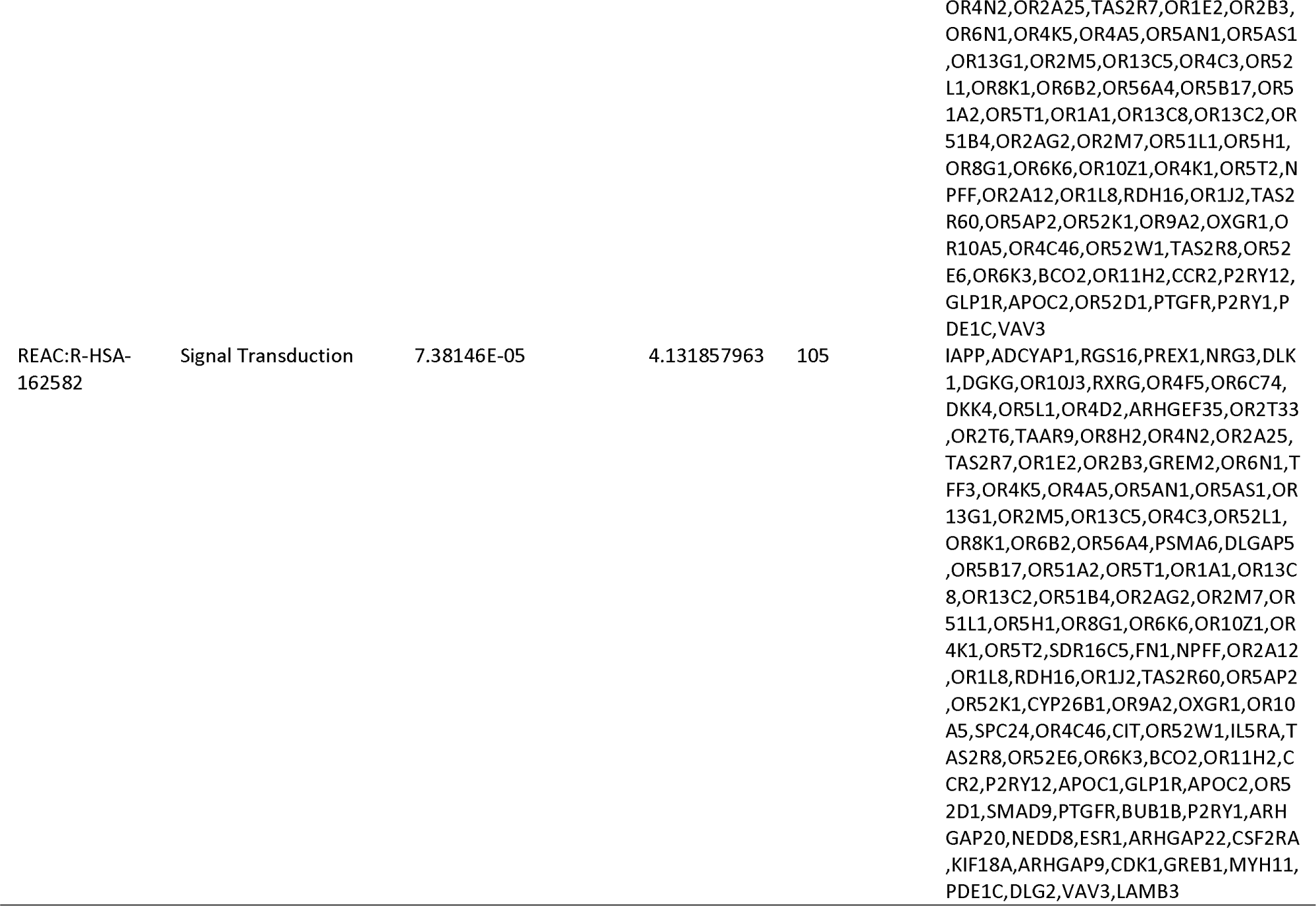
The enriched pathway terms of the up and down regulated differentially expressed genes

### Construction of the PPI network and module analysis

The PPI network of the DEGs was constructed with 5111 nodes and 9392 edges by using the IntAct database (Fig. 3). A node with a higher node degree, betweenness centrality, stress centrality and closeness centrality consider as a hub genes and are listed in Table 5. The key points of the network include the up regulated hub genes, such as MYC, EGFR, LNX1, YBX1 and HSP90AA1, and the down regulated hub genes, such as ESR1, FN1, TK1, ANLN and SMAD9. To detect significant modules in the PPI network, the PEWCC1 plug in was used for analysis, and two modules that had the highest degree stood L out. GO and pathway enrichment analysis showed that module 1 contained 28 nodes and 63 edges (Fig.4A), which were associated with diseases of signal transduction by growth factor receptors and second messengers, disease, nitrogen compound metabolic process and membrane-enclosed lumen, while module 2 had 14 nodes and 30 edges (Fig.4B), which were mainly associated with signal transduction, multicellular organismal process and detection of stimulus.

**Fig. 3.**
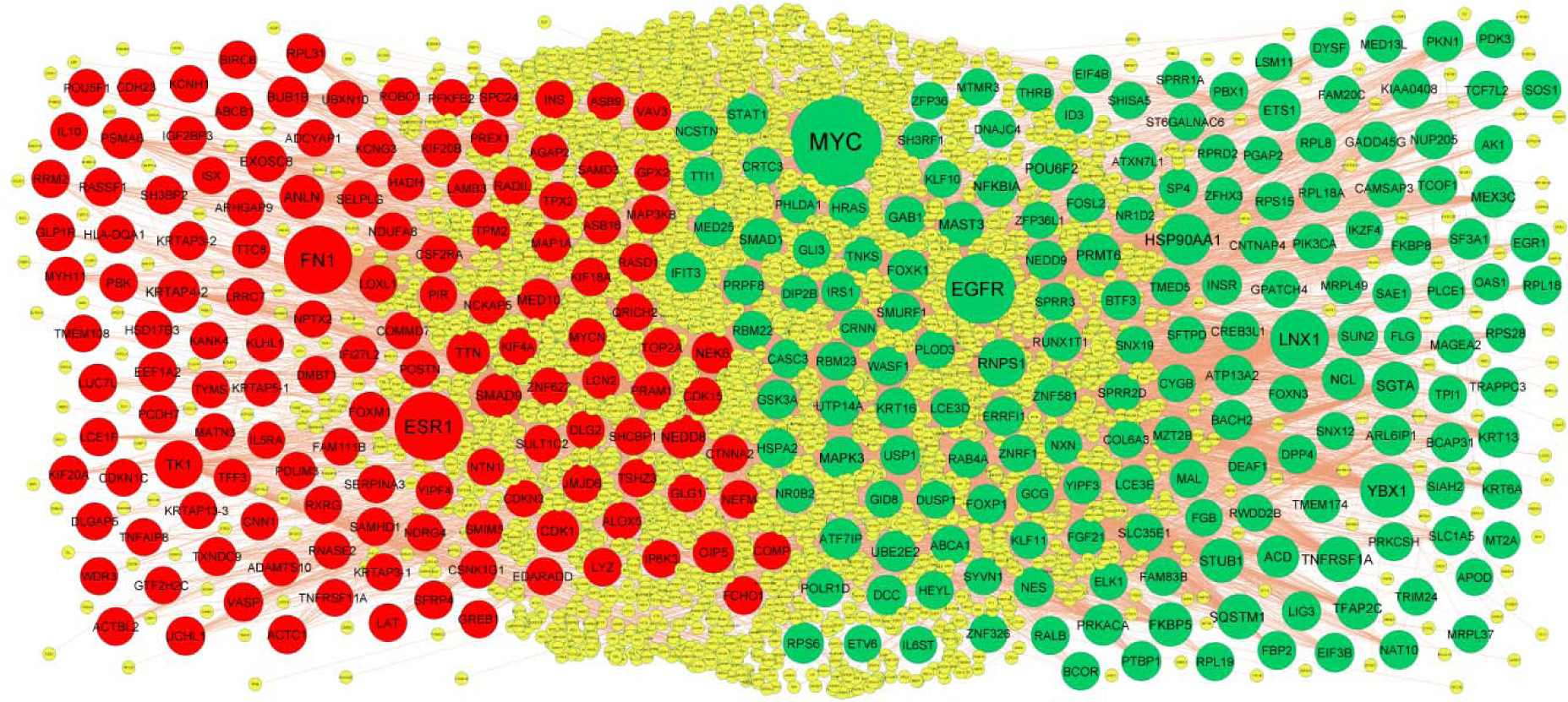
PPI network of DEGs. The PPI network of DEGs was constructed using Cytoscap. Up regulated genes are marked in green; down regulated genes are marked in red

**Fig. 4.**
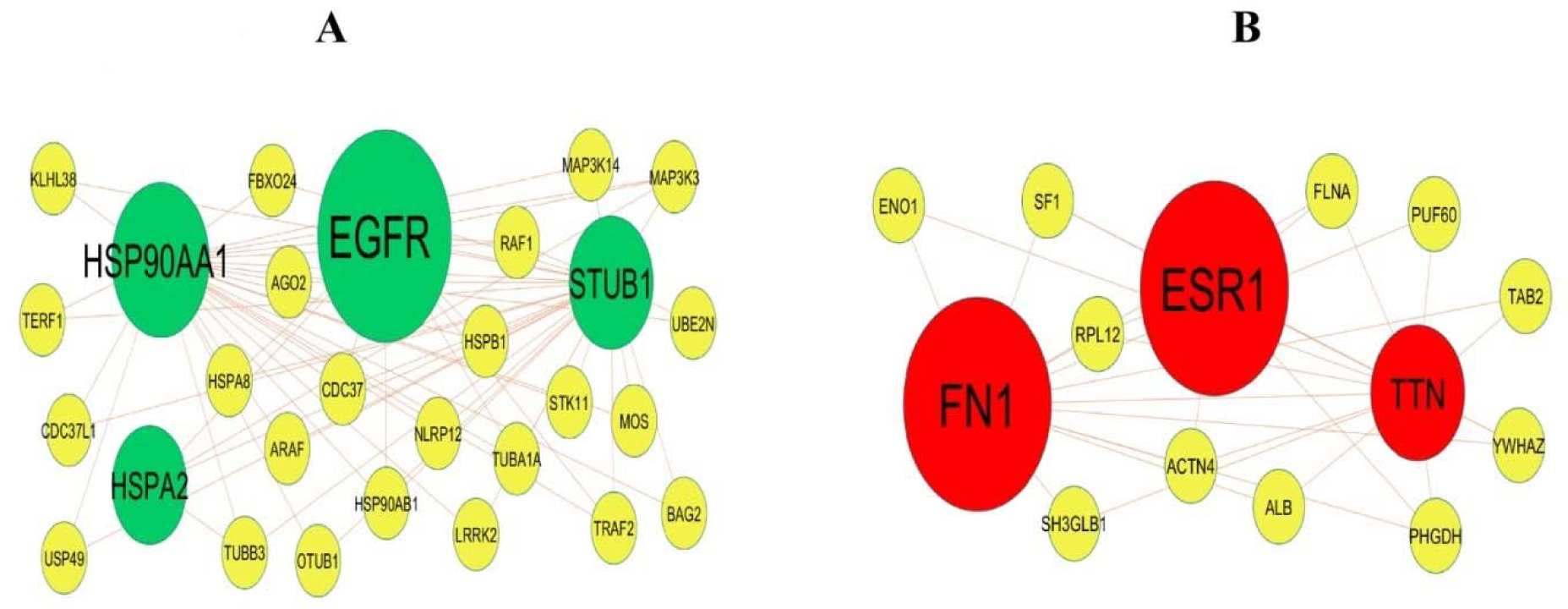
Modules of isolated form PPI of DEGs. (A) The most significant module was obtained from PPI networ with 28 nodes and 63 edges for up regulated genes (B) The most significant module was obtained from PPI network with 14 nodes and 30 edges for down regulated genes. Up regulated genes are marked in green; down regulated genes are marked in red.

**Table 5.**
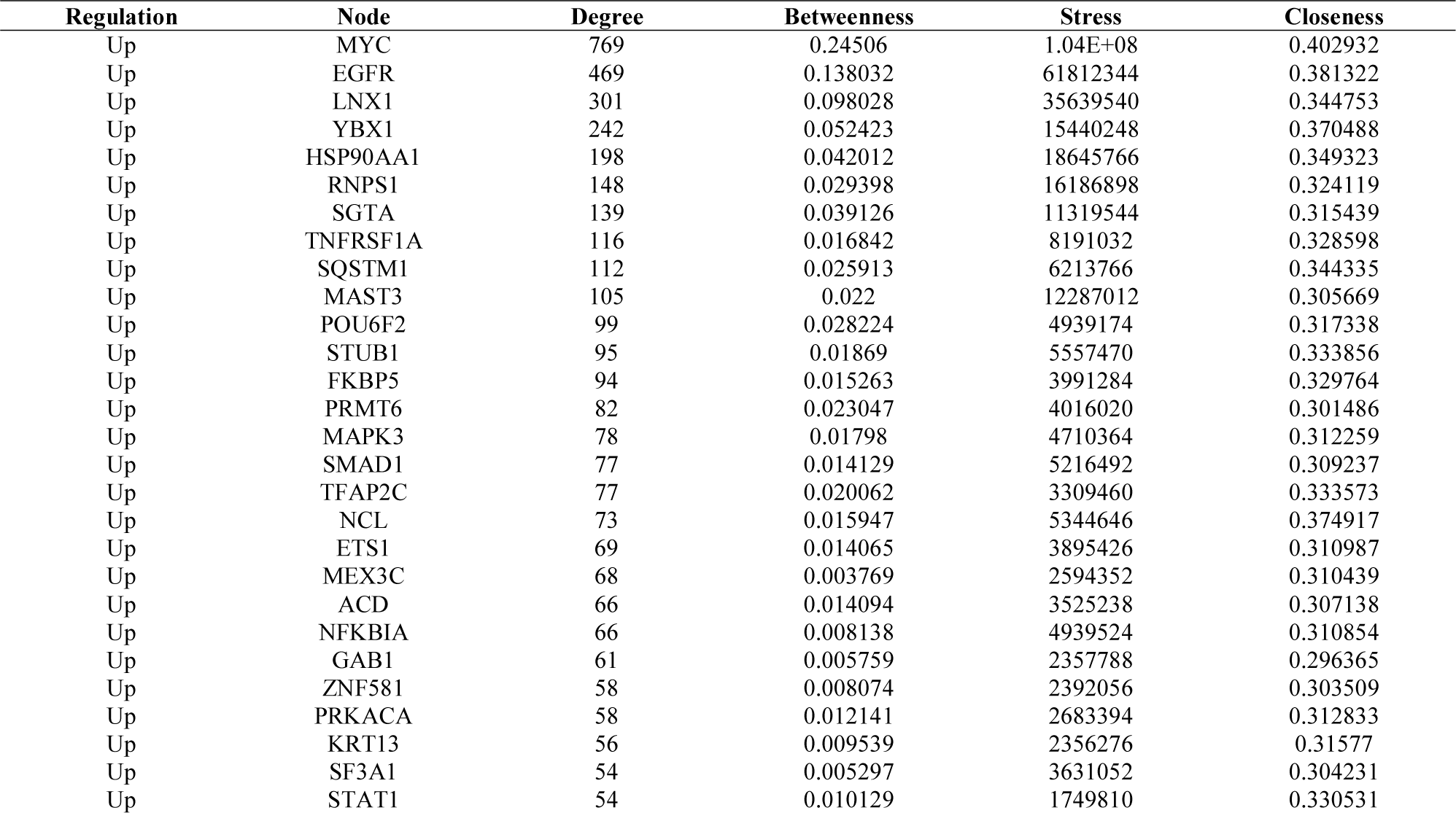
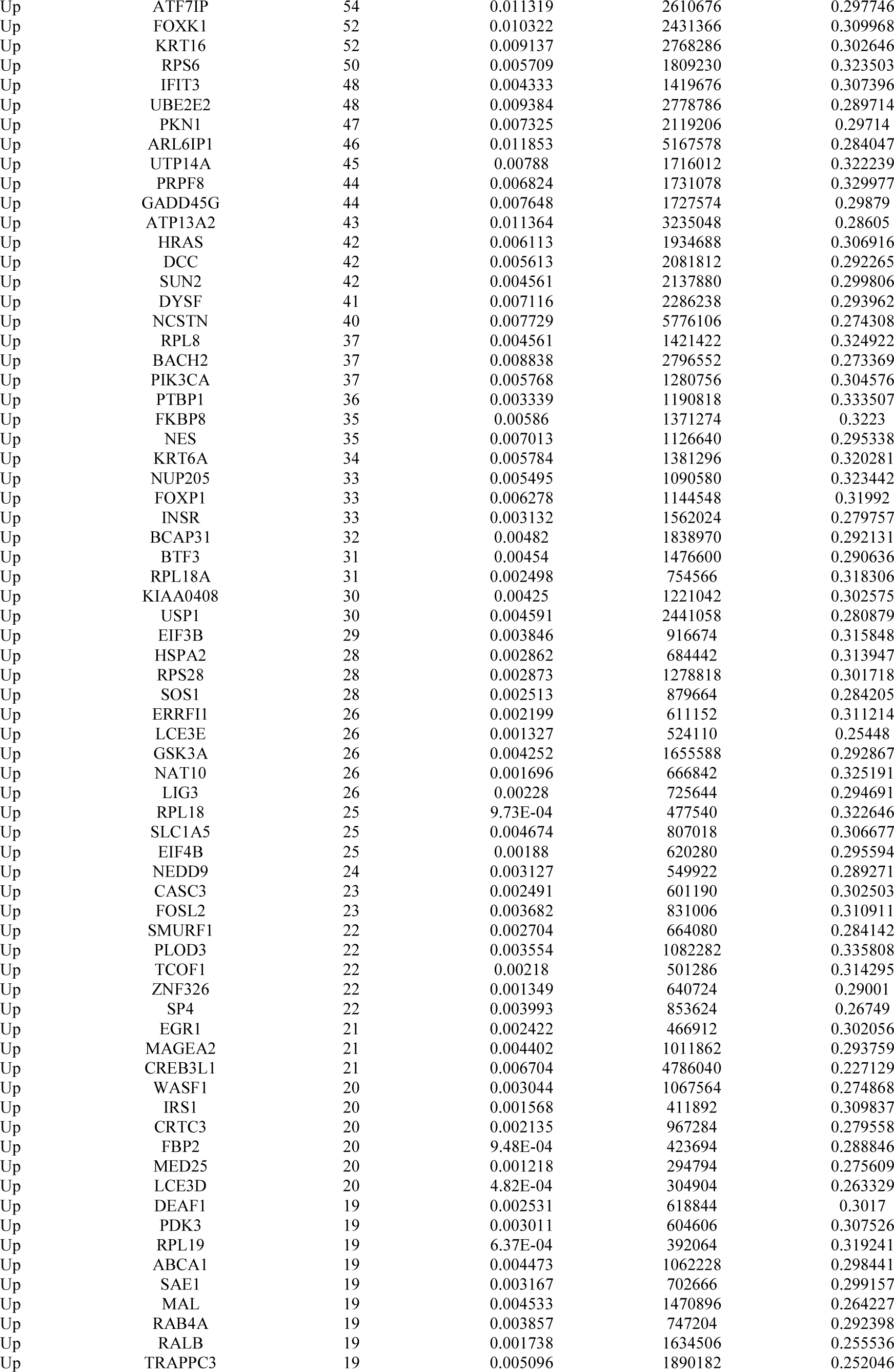
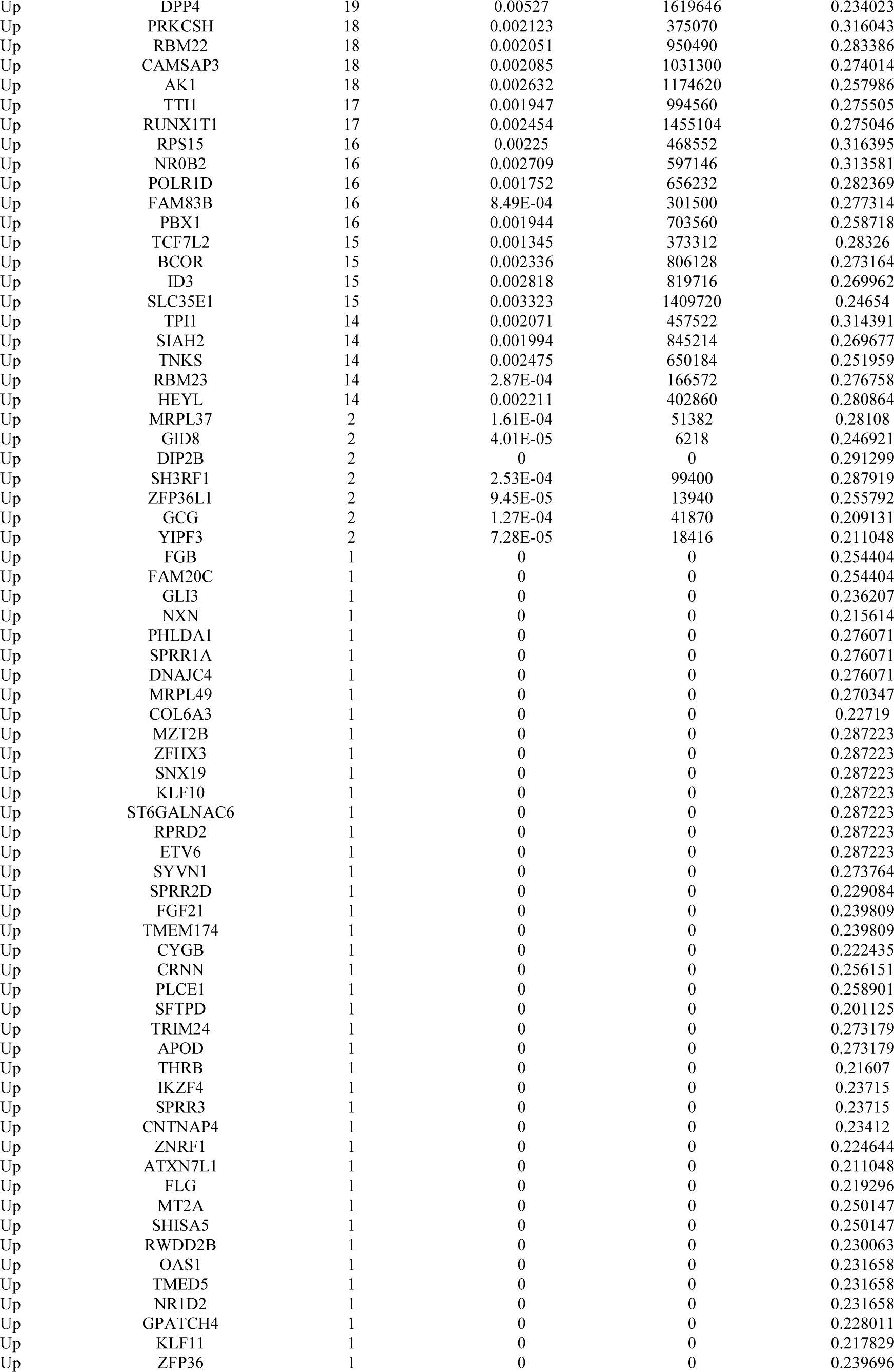
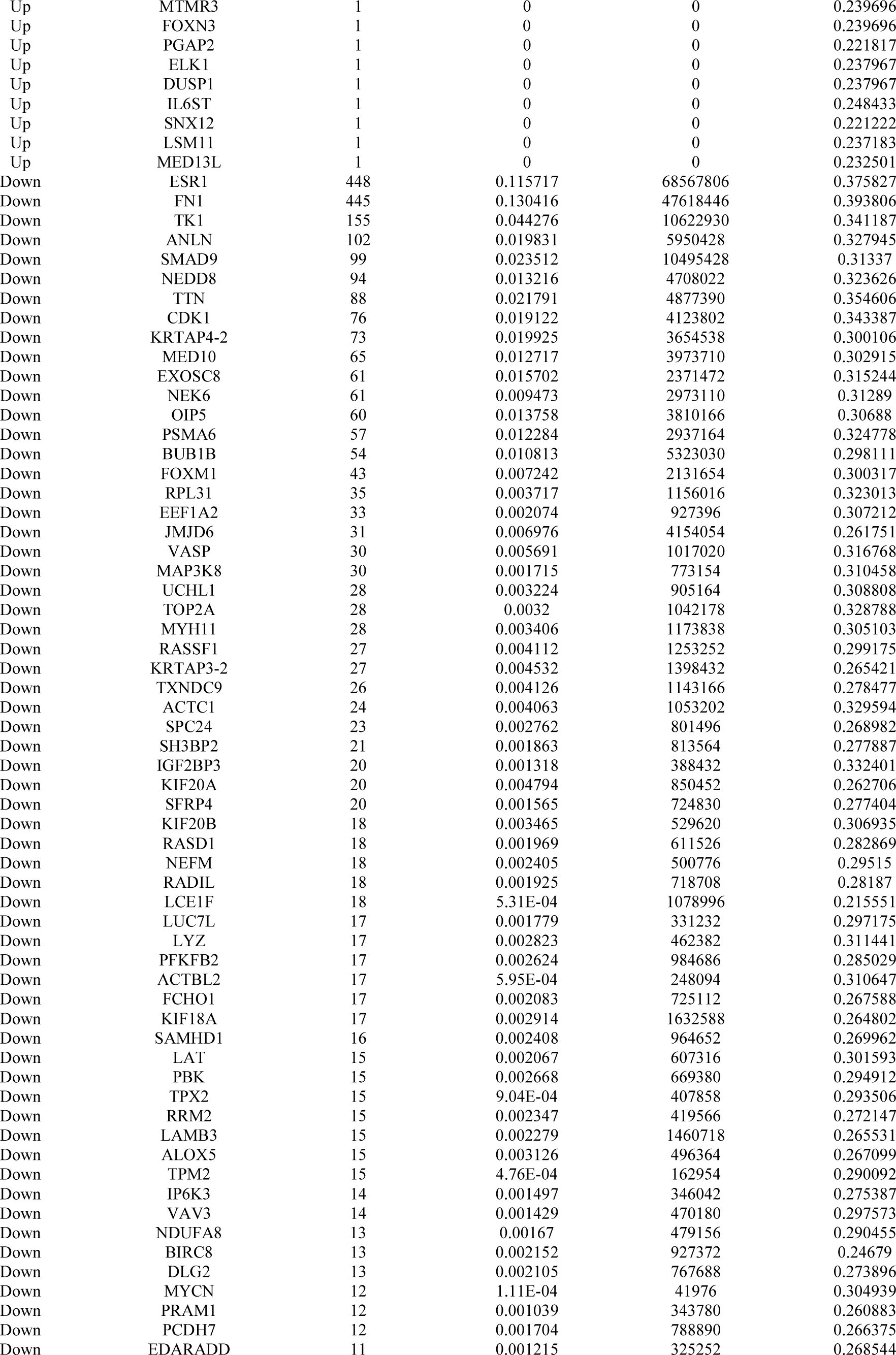
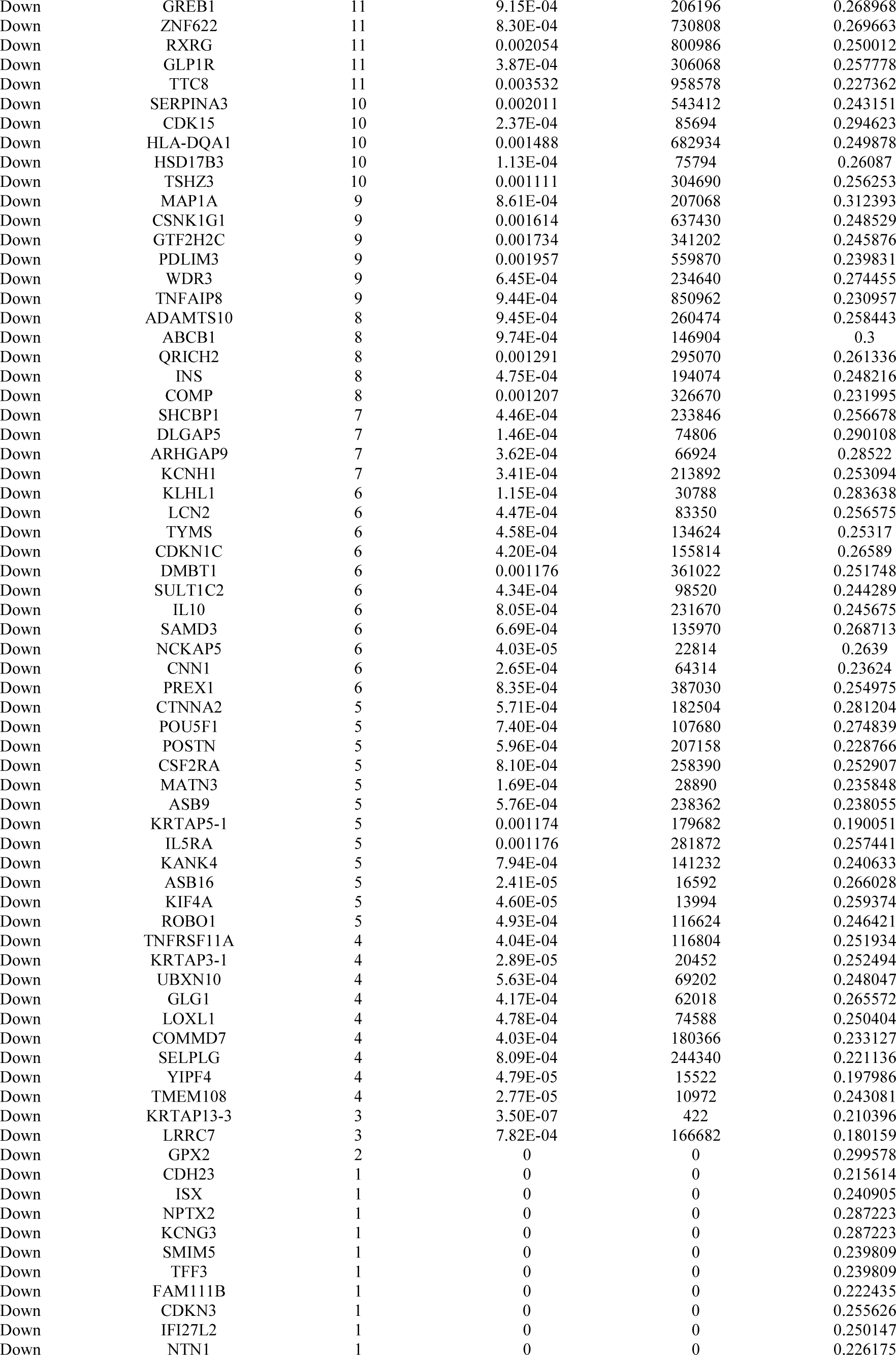
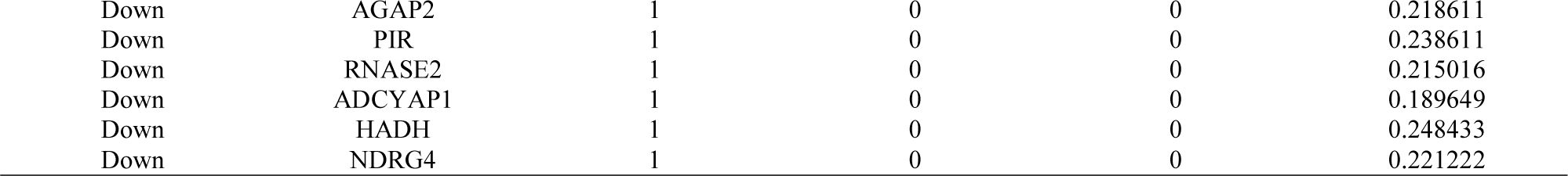
Topology table for up and down regulated genes.

### MiRNA-hub gene regulatory network construction

The network of miRNAs and predicted targets (hub genes) is presented in Table 6. Based on the miRNAs, a miRNA -hub gene regulatory network was constructed with 2568 nodes (miRNA: 2259; hub gene: 309) and 16618 interaction pairs (Fig.5). Notably, MYC targeted 194 miRNAs, including hsa-mir-4677-3p; HSP90AA1 targeted 188 miRNAs, including hsa-mir-3125; FKBP5 targeted 116 miRNAs, including hsa-mir-4779; RNPS1 targeted 109 miRNAs, including hsa-mir-548az-3p; SQSTM1 targeted 108 miRNAs, including hsa-mir-106a-5p; ANLN targeted 127 miRNAs, including hsa-mir-664a-3p; CDK1 targeted 109 miRNAs, including hsa-mir-5688;FN1 targeted 105 miRNAs, including hsa-mir-199b-3p;ESR1 targeted 98 miRNAs, including hsa-mir-206; TK1 targeted 80 miRNAs, including hsa-mir-6512-3p.

**Fig. 5.**
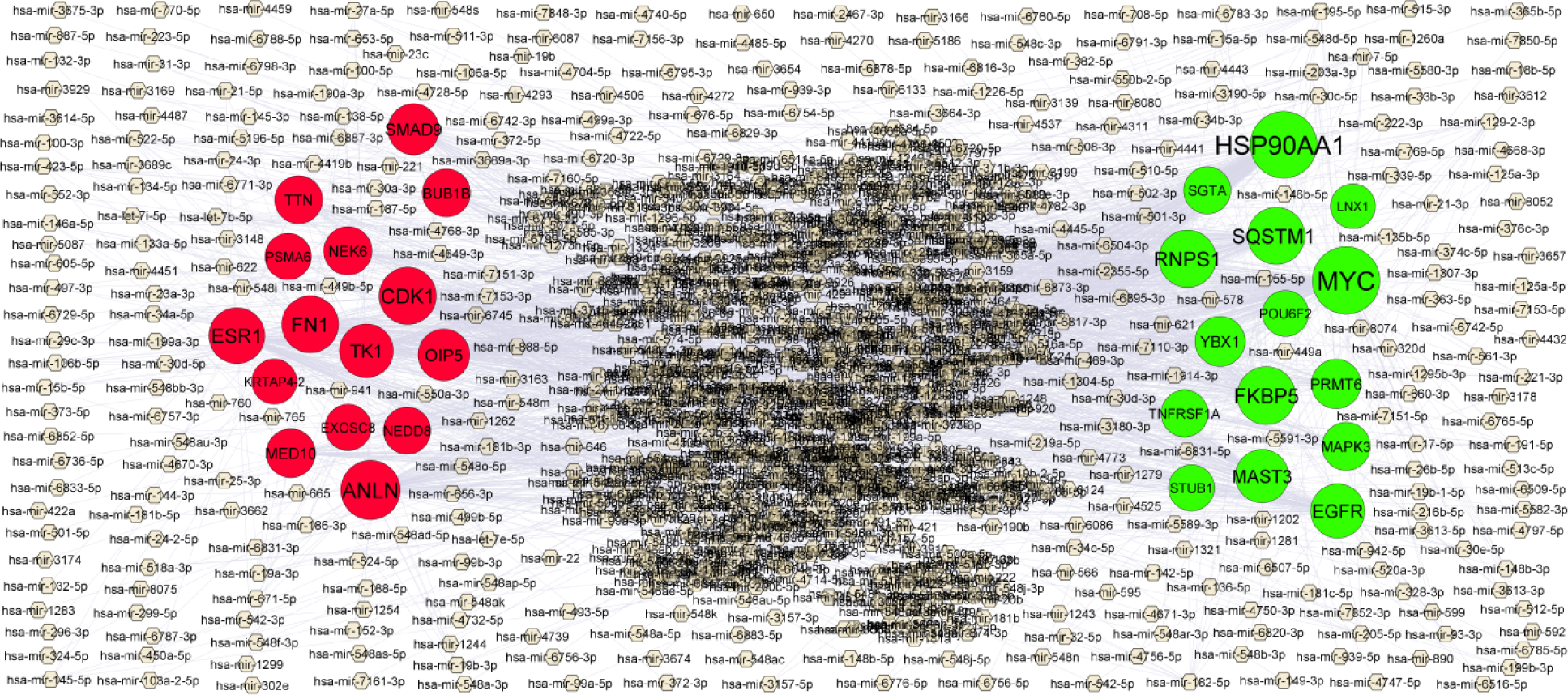
MiRNA - hub gene regulatory network. The chocolate color diamond nodes represent the key miRNAs; up regulated genes are marked in green; down regulated genes are marked in red.

**Table 6.** miRNA - target gene and TF - target gene interaction

| Regulation | Target Genes | Degree | MicroRNA | Regulation | Target Genes | Degree | TF |
| --- | --- | --- | --- | --- | --- | --- | --- |
| Up | MYC | 194 | hsa-mir-4677-3p | Up | MAPK3 | 48 | JUND |
| Up | HSP90AA1 | 188 | hsa-mir-3125 | Up | HSP90AA1 | 35 | HSF2 |
| Up | FKBP5 | 116 | hsa-mir-4779 | Up | SQSTM1 | 34 | SMAD4 |
| Up | RNPS1 | 109 | hsa-mir-548az-3p | Up | STUB1 | 31 | ATF6 |
| Up | SQSTM1 | 108 | hsa-mir-106a-5p | Up | EGFR | 27 | ELF3 |
| Up | EGFR | 83 | hsa-mir-219a-5p | Up | YBX1 | 24 | TFAP2A |
| Up | MAST3 | 76 | hsa-mir-129-2-3p | Up | LNK1 | 21 | MAX |
| Up | YBX1 | 48 | hsa-mir-1537-3p | Up | FKBP5 | 20 | PGR |
| Up | PRMT6 | 43 | hsa-mir-4330 | Up | PRMT6 | 9 | YY1 |
| Up | MAPK3 | 33 | hsa-mir-3158-3p | Up | SGTA | 7 | MXI1 |
| Up | SGTA | 24 | hsa-mir-421 | Up | POU6F2 | 7 | ALX1 |
| Up | TNFRSF1A | 22 | hsa-mir-548an | Up | RNPS1 | 7 | STAT3 |
| Up | STUB1 | 16 | hsa-mir-942-5p | Up | TNFRSF1A | 6 | EP300 |
| Up | POU6F2 | 15 | hsa-mir-7850-5p | Up | MAST3 | 1 | NFYA |
| Down | ANLN | 127 | hsa-mir-664a-3p | Down | ESR1 | 126 | FOXF2 |
| Down | CDK1 | 109 | hsa-mir-5688 | Down | SMAD9 | 38 | XAB2 |
| Down | FN1 | 105 | hsa-mir-199b-3p | Down | CDK1 | 36 | KHDRBS1 |
| Down | ESR1 | 98 | hsa-mir-206 | Down | FN1 | 25 | RELA |
| Down | TK1 | 80 | hsa-mir-6512-3p | Down | NEK6 | 16 | SRY |
| Down | OIP5 | 62 | hsa-mir-767-5p | Down | TK1 | 11 | DRAP1 |
| Down | SMAD9 | 58 | hsa-mir-3689a-3p | Down | NEDD8 | 10 | PARP1 |
| Down | MED10 | 41 | hsa-mir-647 | Down | TTN | 10 | FOXD3 |
| Down | NEK6 | 34 | hsa-mir-4485-3p | Down | ANLN | 9 | JUN |
| Down | BUB1B | 32 | hsa-mir-449b-5p | Down | BUB1B | 9 | MYB |
| Down | TTN | 31 | hsa-mir-181c-5p | Down | MED10 | 6 | KDM4B |
| Down | NEDD8 | 26 | hsa-mir-583 | Down | PSMA6 | 6 | EBF1 |
| Down | PSMA6 | 17 | hsa-mir-539-5p | Down | OIP5 | 5 | GATA2 |
| Down | EXOSC8 | 17 | hsa-mir-191-5p | Down | KRTAP4-2 | 1 | TBP |

### TF-hub gene regulatory network construction

The network of TFs and predicted targets (hub genes) is presented in Table 6. Based on the TFs, a TF -hub gene regulatory network was constructed with 899 nodes (TF: 604; hub gene: 295) and 3542 interaction pairs (Fig.6). Notably, MAPK3 targeted 48 TFs, including JUND; HSP90AA1 targeted 35 TFs, including HSF2; SQSTM1 targeted 34 TFs, including SMAD4; STUB1 targeted 31 TFs, including ATF6; EGFR targeted 27 TFs, including ELF3; ESR1 targeted 126 TFs, including ELF3; SMAD9 targeted 38 TFs, including ELF3; CDK1 targeted 36 TFs, including ELF3; FN1 targeted 25 TFs, including ELF3; NEK6 targeted 16 TFs, including ELF3.

**Fig. 6.**
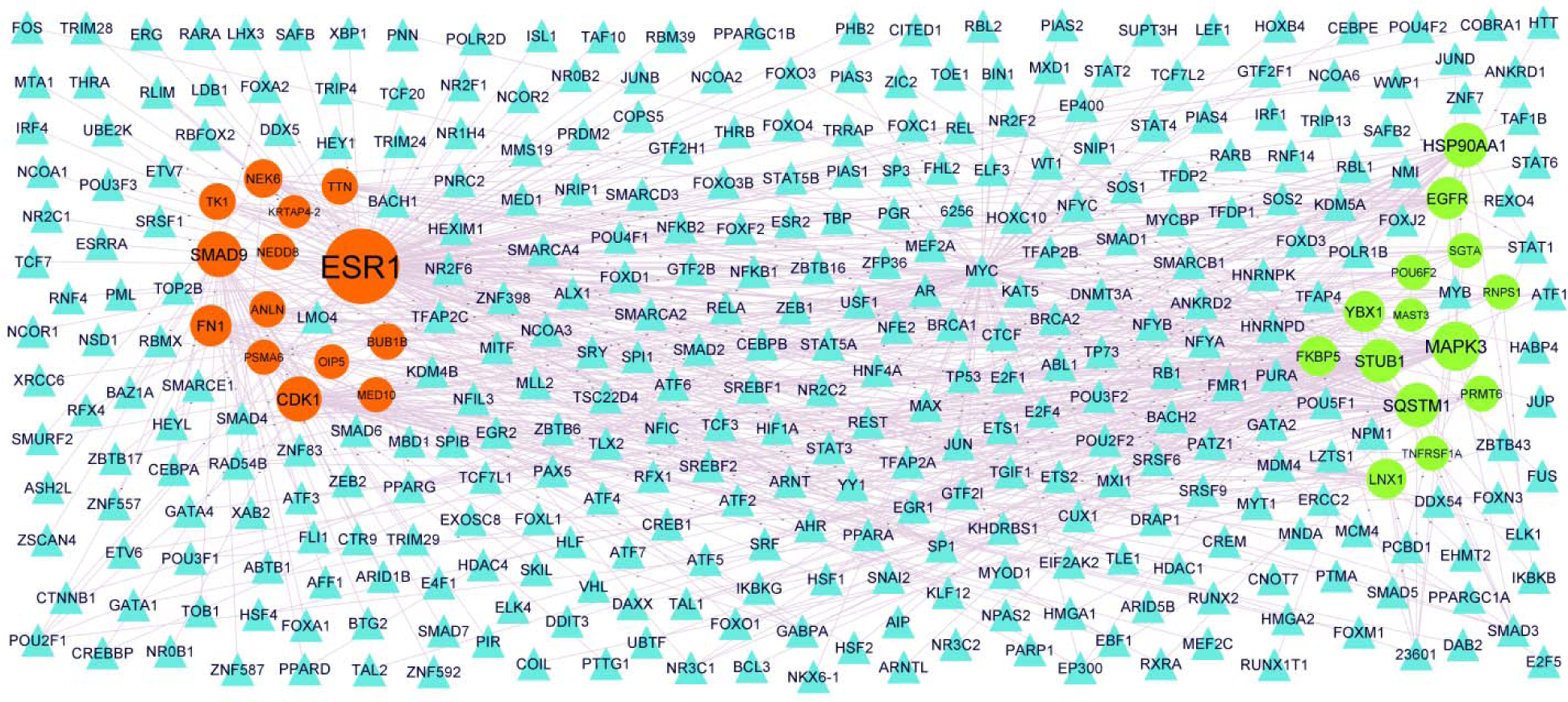
TF - hub gene regulatory network. The blue color triangle nodes represent the key TFs; up regulated genes are marked in green; down regulated genes are marked in red.

### Validation of hub genes by receiver operating characteristic curve (ROC) analysis

ROC curve analysis was for evaluating the capacity of MYC, EGFR, LNX1, YBX1, HSP90AA1, ESR1, FN1, TK1, ANLN and SMAD9, so as to distinguish T1DM from normal control (Fig.7). ROC curve analysis showed that MYC, EGFR, LNX1, YBX1, HSP90AA1, ESR1, FN1, TK1, ANLN and SMAD9 expression levels have potential diagnostic value for T1DM patients (AUCL>L0.9).

**Fig. 7.**
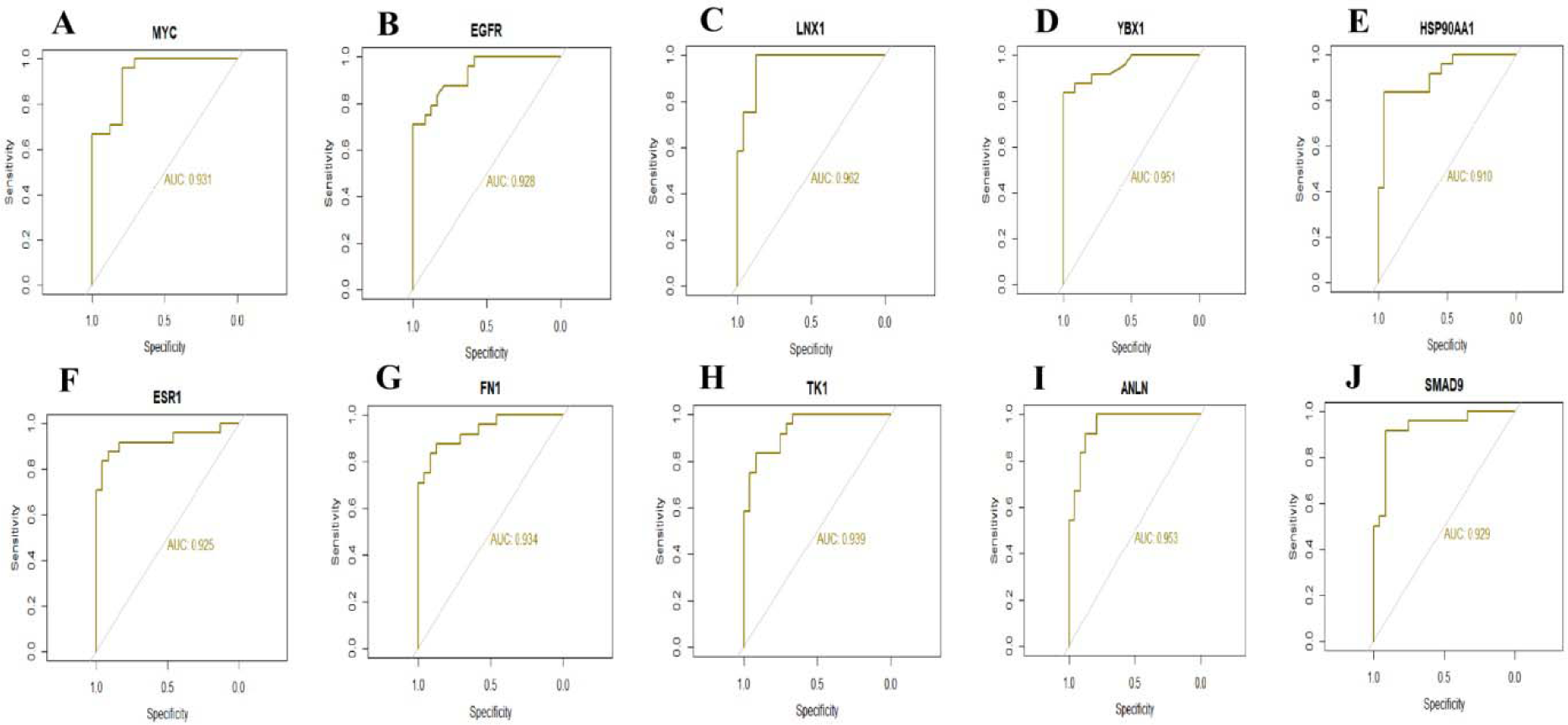
ROC curve validated the sensitivity, specificity of hub genes as a predictive biomarker for dementia prognosis. A) MYC B) EGFR C) LNX1 D) YBX1 E) HSP90AA1 F) ESR1 G) FN1 H) TK1 I) ANLN J) SMAD9

### Detection of the mRNA expression of the hub genes by RT-PCR

Next, in order to verify the results of previous bioinformatics analysis, the gene expression levels of MYC, EGFR, LNX1, YBX1, HSP90AA1, ESR1, FN1, TK1, ANLN and SMAD9 were detected by RT-PCR between T1DM and normal control. As shown in Fig 8, compared with normal tissues, MYC, EGFR, LNX1, YBX1 and HSP90AA1 mRNA expression levels were significantly up regulated in the T1DM, and ESR1, FN1, TK1, ANLN and SMAD9 mRNA level were down regulated, which was consistent with the results of bioinformatics analysis.

**Fig. 8.**
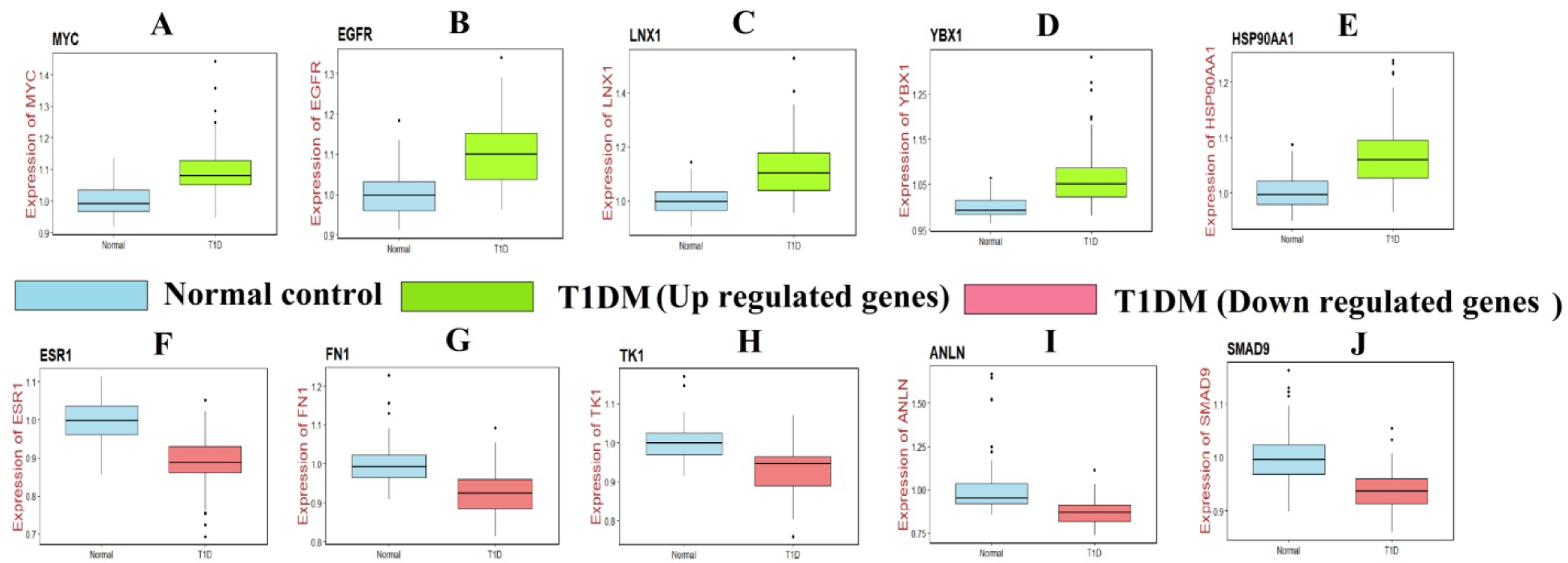
Validation of hub genes by RT- PCR. A) MYC B) EGFR C) LNX1 D) YBX1 E) HSP90AA1 F) ESR1 G) FN1 H) TK1 I) ANLN J) SMAD9

## Discussion

T1DM is the common forms of chronic autoimmune diabetes that affect an individual’s quality of life [43]. However, the potential causes of T1DM remain uncertain. Understanding the underlying molecular pathogenesis of T1DM is of key importance for diagnosis, prognosis, and identifying drug targets. As high-throughput RNA sequencing can provide information regarding the expression levels of thousands of genes in the human genome simultaneously, this methodology has been widely used to predict the potential diagnostic and therapeutic targets for T1DM. In the present investigation, we extracted the data from GSE162689, which includes 27 T1DM samples and 32 normal control samples. We identified 477 up regulated and 475 down regulated genes between T1DM samples and normal control samples using bioinformatics analysis. FGA (fibrinogen alpha chain) [44] and FGB (fibrinogen beta chain) [45] genes have been found to be expressed in cardiovascular disease, but these genes might be novel target for T1DM. IGF2 [46], IAPP (islet amyloid polypeptide) [47], INS (insulin) [48] and MAFA (MAF bZIP transcription factor A) [49] have been reported be associated with the development of T1DM. Previous study have indicated the expression of ADCYAP1 is associated in type 2 diabetes mellitus [50], but this gene might be novel target for T1DM. Gold et al. [51] found the CSNK1G1 was a prognostic factor in cognitive impairment, but this gene might be novel target for T1DM.

Furthermore, we investigated the biological functions of these DEGs by using online website, and GO and pathway enrichment analysis. Husemoen et al [52], Zhang et al [53], Hartz et al [54], Słomiń ki et al [55], Johansson et al [56], sPan et al [57], Lopez-Sanz et al [58], Grant, [59], Słomiń i et al [60], Galán et al sk [61], Jordan et al [62], Winkler et al [63], Yip et al [64], Crookshank et al [65], Lempainen et al [66], Qu and Polychronakos, [67], Morrison et al [68], Zhang et al [69], Gerlinger-Romero et al [70], Belanger et al [71], Dieter et al [72], Wanic et al [73], Ushijima Wanic et al [74], Guo et al [75], Davis et al [76], Elbarbary et al [77], Villasenor et al [78], Zhang et al [79], Lee et al [80], Zhi et al [81], Li Calzi et al [82], Sebastiani et al [83], Cherney et al [84], Doggrell, [85] and Yanagihara et al [86] found that FLG (filaggrin), FGF21, PEMT (phosphatidylethanolamine N-methyltransferase) KL (klotho), CEL (carboxyl ester lipase), FOSL2, STAT1, TCF7L2, TP53, EGFR (epidermal growth factor receptor), ETS1, KCNJ8, DEAF1, GCG (glucagon), IKZF4, OAS1, IRS1, ABCG2, FBXO32, PTBP1, BACH2, CNDP2, KLF11, MT1E, DPP4, SLC29A3, RGS16, MAS1, GCGR (glucagon receptor), HLA-C, VASP (vasodilator stimulated phosphoprotein), CCR2, PTGS2, GLP1R and JMJD6 are involved in the progression of T1DM. Vassilev et al [87], Qin et al [88], Ma et al [89], West et al [90], Hoffmann et al [91], Deary et al [92], Belangero et al [93], Jung et al [94], Tang et al [95], Goodier et al [96], Petyuk et al [97], Roux et al [98], Castrogiovanni et al [99], Suleiman et al [100], Haack et al [101], Kwiatkowski et al [102], Pinacho et al [103], Luo et al [104], He et al [105], Moudi et al [106], Thevenon et al [107], Li et al [108], Reitz et al [109], Jenkins and Escayg [110], Letronne et al [111], Ma et al [112], Chabbert et al [113], Abramsson et al [114], Aeby et al [115] and Roll et al [116] found that the expression of DCC (DCC netrin 1 receptor), PLP1, SNX19, SH3RF1, TNFRSF1A, NCSTN (nicastrin), DGCR2, NPAS2, CDNF (cerebral dopamine neurotrophic factor), SMCR8, HSPA2, STUB1, CHID1, ATP13A2, SQSTM1, LIG3, SP4, ACSL6, ERN1, ATF6B, LRFN2, NRG3, LRRTM3, GABRA2, ADAM30,

GABRR2, TSHZ3, LOXL1, SCN1B and SRPX2 are associated with the prognosis of patients with cognitive impairment, but these genes might be novel target for T1DM. Recent studies found that KCP (kielin cysteine rich BMP regulator) [117], NOG (noggin) [118], COL6A3 [119], BTG2 [120], RPS6 [121], KLF15 [122], KLF3 [123], ZFP36 [124], ETV5 [125], TLE3 [126], NNMT (nicotinamide N-methyltransferase) [127], WDTC1 [128], ZFHX3 [129], SIAH2 [130], MBOAT7 [131], RUNX1T1 [132], MAPK4 [133], KLF9 [134], SELENBP1 [135], HELZ2 [136], ELK1 [137], SERTAD2 [138], CRTC3 [139], ABCB11 [140], TACR1 [141], SLC22A11 [142], PER3 [143], P2RX5 [144], MFAP5 [145], FGL1 [146], OLFM4 [147], NTN1 [148], ESR1 [149], ABCB1 [150], VAV3 [151] and LAMB3 [152] plays an important role in the occurrence and development of obesity, but these genes might be novel target for T1DM. STAR (steroidogenic acute regulatory protein) [153], IL1RN [154], AQP5 [155], EGR1 [156], SFTPD (surfactant protein D) [157], KLF10 [158], PODXL (podocalyxin like) [159], FOXN3 [160], IL6R [161], PBX1 [162], APOD (apolipoprotein D) [163], ACVR2B [164], CD34 [165], INSR (insulin receptor) [166], APOA5 [167], STAR (steroidogenic acute regulatory protein) [168], PDK4 [169], GLS (glutaminase) [170], FKBP5 [171], SLC6A15 [172], MT2A [173], SLC38A4 [174], AQP7 [175], ABHD15 [176], ABCA1 [177], ZNRF1 [178], PPP1R3B [179], MAOA (monoamine oxidase A) [180], UBE2E2 [181], RNASEK (ribonuclease K) [182], PREX1 [183], DGKG (diacylglycerol kinase gamma) [184], POSTN (periostin) [185], COMP (cartilage oligomeric matrix protein) [186], GAP43 [187], P2RY12 [188], SELL (selectin L) [189] and DLG2 [190] have been revealed to be associated with type 2 diabetes mellitus, but these genes might be novel target for T1DM. Expression of ERRFI1 [191], ALOX12 [192], SOCS5 [193], DDIT4 [194], DUSP4 [195], IL6ST [196], DUSP1 [197], SMAD1 [198], NCL (nucleolin) [199], METTL14 [200], FMOD (fibromodulin) [201], CYGB (cytoglobin) [202], UNC5A [203] and TAAR9 [204] are associated with prognosis in patients with diabetic nephropathy, but these genes might be novel target for T1DM. A previous study reported that FAP (fibroblast activation protein alpha) [205], EYA4 [206], BCL9 [207], IRF2BP2 [208], EGR3 [209], GADD45B [210], DMD (dystrophin) [211], LSR (lipolysis stimulated lipoprotein receptor) [212], DLL4 [213], SUN2 [214], SOS1 [215], PIK3CA [216], GAMT (guanidinoacetate N-methyltransferase) [217], RBM47 [218], HSP90AA1 [219], GAB1 [220], S1PR1 [221], EDNRB (endothelin receptor type B) [222], NFKBIA (NFKB inhibitor alpha) [223], GJA1 [224], GADD45G [225], PHLDA1 [226], CMPK2 [227], FIGN (fidgetin, microtubule severing factor) [228], KCNJ2 [229], ABCC9 [230], DIRAS3 [231], EPHX1 [232], RAB4A [233], UBIAD1 [234], CASQ2 [235], TTN (titin) [236], KCNH1 [237], JPH2 [238], OXGR1 [239], UCHL1 [240], SERPINA3 [241], MMP28 [242], ADAMTS2 [243], P2RY1 [244], CSF2RA [245], MYO1F [246], SELPLG (selectin P ligand) [247] and SAMHD1 [248] are expressed in cardiovascular disease, but these genes might be novel target for T1DM. MAOB (monoamine oxidase B) [249], VEGFC (vascular endothelial growth factor C) [250], DBP (D-box binding PAR bZIP transcription factor) [251], MYADM (myeloid associated differentiation marker) [252], NES (nestin) [253], SMURF1 [254], EDNRB (endothelin receptor type B) [255], MUC6 [256], TOR2A [257], TNKS (tankyrase) [258], NEDD9 [259], ASIC1 [260], ADAMTS8 [261], DYSF (dysferlin) [262], SLC26A9 [263], SLC45A3 [264] and KCNQ2 [265] contributes to the progression of hypertension, but these genes might be novel target for T1DM. Yang et al [266], Zhang et al [267] and Wang et al [268] demonstrated that SYVN1, BTG1 and CFB (complement factor B) were associated with diabetic retinopathy, but these genes might be novel target for T1DM.

PPI network and modules were used to identify hub genes. The results of the present investigation might provide potential biomarkers for the diagnosis of T1DM. SMAD9 has been shown to be activated in hypertension [269], but this genes might be novel target for T1DM. MYC, LNX1, YBX1, FN1, TK1 and ANLN (anillin actin binding protein) might be the novel biomarkers for T1DM.

In this investigation, the miRNA-hub gene regulatory network and TF-hub gene regulatory network that regulates T1DM was constructed. Recent studies found that CDK1 [270], hsa-mir-199b-3p [271], JUND [272] and FOXF2 [273] plays an important role in the occurrence and development of obesity, but these genes might be novel target for T1DM. The hsa-mir-106a-5p [274], hsa-mir-206 [275], SMAD4 [276] and ATF6 [277] are a major regulator of type 2 diabetes mellitus, but these genes might be novel target for T1DM. Studies have shown that the hsa-mir-106a-5p [278] and HSF2 [279] are essential for regulating cardiovascular disease, but these genes might be novel target for T1DM. Mendes-Silva et al [280] indicated that hsa-mir-664a-3p facilitated cognitive impairment, but this gene might be novel target for T1DM. ELF3 participated in diabetic nephropathy [281], but this gene might be novel target for T1DM. SRY is crucial for hypertension progression [282], but this gene might be novel target for T1DM. RNPS1, MAPK3, NEK6, hsa-mir-4677-3p, hsa-mir-3125, hsa-mir-4779, hsa-mir-548az-3p, hsa-mir-5688, hsa-mir-6512-3p, XAB2, KHDRBS1 and RELA might be the novel biomarkers for T1DM.

In conclusion, the present investigation shows the global profile of DEGs and relative signaling pathways that might play in the initiation and progression of T1DM. In the pathogenesis of acne, the possible crucial genes are MYC, EGFR, LNX1, YBX1, HSP90AA1, ESR1, FN1, TK1, ANLN and SMAD9, and the possible important GO terms and pathways are multicellular organism development, detection of stimulus, diseases of signal transduction by growth factor receptors and second messengers, and olfactory signaling pathway. Therefore, it provides new research directions for the detection and treatment of T1DM. However, their involvement in the molecular mechanisms of disease needs further clinical studies.

## Acknowledgement

I thank Anders Roger Hedin, Uppsala University, Clinical immunology, Olle Korsgren, Uppsala, Uppland, Sweden, very much, the author who deposited their profiling by high throughput sequencing dataset GSE162689, into the public GEO database.

## Ethical approval

This article does not contain any studies with human participants or animals performed by any of the authors.

## Informed consent

No informed consent because this study does not contain human or animals participants.

## Availability of data and materials

The datasets supporting the conclusions of this article are available in the GEO (Gene Expression Omnibus) (https://www.ncbi.nlm.nih.gov/geo/) repository. [(GSE162689) (https://www.ncbi.nlm.nih.gov/geo/query/acc.cgi?acc=GSE162689]

## Consent for publication

Not applicable.

## Author Contributions

B. V - Writing original draft, and review and editing

C. V - Software and investigation

## Authors

Basavaraj Vastrad ORCID ID: 0000-0003-2202-7637 Chanabasayya Vastrad ORCID ID: 0000-0003-3615-4450

